# The longevity effects of reduced IGF-1 signaling depend on the stability of the mitochondrial genome

**DOI:** 10.1101/2025.06.03.656903

**Authors:** Sarah J. Shemtov, Eric McGann, Lucy Carrillo, Sangmin Lee, Herbert Anson, Claire S. Chung, Maria-Eleni Anagnostou, Guan-Ju D. Lai, Bert M. Verheijen, Junxiang Wan, Ivetta Vorobyova, Monica Sanchez-Contreras, Cheryl A. Conover, Max A. Thorwald, Pinchas Cohen, Scott R. Kennedy, Jean-François Gout, Suraiya Haroon, Marc Vermulst

## Abstract

Suppression of insulin-like growth factor-1 (IGF-1) signaling extends mammalian lifespan and protects against a range of age-related diseases. Surprisingly though, we found that reduced IGF-1 signaling fails to extend the lifespan of mitochondrial mutator mice. Accordingly, most of the longevity pathways that are normally initiated by IGF-1 suppression were either blocked or blunted in the mutator mice. These observations suggest that the pro-longevity effects of IGF-1 suppression critically depend on the integrity of the mitochondrial genome and that mitochondrial mutations may impose a hard limit on mammalian lifespan. Together, these findings deepen our understanding of the interactions between the hallmarks of aging and underscore the need for interventions that preserve the integrity of the mitochondrial genome.

## INTRODUCTION

Aging is a complex biological process that is characterized by multiple molecular and cellular hallmarks, including genomic instability, mitochondrial dysfunction and deregulated nutrient sensing^1^. While significant progress has been made in our understanding of the individual hallmarks of human aging, their interactions remain poorly defined. In addition to obscuring our insight into the basic biology of aging, this gap in our knowledge complicates the development of effective anti-aging interventions, as targeting one hallmark of aging may inadvertently exacerbate others.

One strategy to elucidate the relationships between the hallmarks of aging is to investigate how the disruption of one hallmark affects the trajectory of another. In doing so, it may be possible to assess whether these processes act independently, synergistically, or in opposition of each other as they shape human lifespan. In addition, this strategy may reveal if a hierarchy exists between aging pathways, which could lead to a more integrated and causally ordered model of the aging process. In this study, we apply this strategy to investigate the relationship between two critical drivers of the aging process, mitochondrial mutagenesis^2^ and IGF-1 signaling^3^.

A large body of evidence supports the idea that instability of the mitochondrial genome (mtDNA) leads to a progressive decline in mitochondrial function, which accelerates the natural aging process and contributes to a wide variety of age-related diseases, including sarcopenia, neurodegeneration and heart failure^2^. A similar body of work describes the role of IGF-1 signaling in the aging process. IGF-1 regulates the growth and metabolism of human tissues, and reduced IGF-1 signaling can not only extend mammalian lifespan, but can also confer resistance against various age-related diseases, including neurodegeneration, metabolic decline, and cardiovascular disease^3^. However, how mitochondrial mutagenesis and IGF-1 signaling interact with each other to shape mammalian lifespan remains unclear.

It is likely though, that mitochondrial mutagenesis and IGF-1 signaling intersect at multiple decisive junctions during the aging process. At a molecular level, mtDNA mutations are a constant source of oxidative stress^4^, a form of cellular damage that is directly influenced by IGF-1 activity^5^. At a systemic level, mitochondria serve as central hubs of metabolic activity and respond dynamically to changes in nutrient availability and growth signals^6^, processes that are tightly regulated by IGF-1 signaling^7^. And finally, at a macroscopic level, mitochondrial mutations directly contribute to a variety of age-related diseases^2,8,9^, many of which can be prevented, or slowed down by reduced IGF-1 signaling^10–14^.

Together, these observations suggest that mitochondrial mutagenesis and IGF-1 signaling do not operate in isolation, but instead converge on overlapping biological pathways that shape the rate and quality of aging. If so, it may be possible to counteract the harmful effects of mtDNA mutations by suppressing IGF-1 signaling, which could improve antioxidant defenses^5^, enhance mitochondrial turnover^15^, and delay the onset of age-related diseases^16^. In support of this idea, we previously found that reduced insulin/IGF-1 signaling can alleviate muscle dysfunction in a *C. elegans* model of mtDNA disease^17^. Alternatively, mitochondrial mutagenesis and IGF-1 signaling may act through parallel, but mechanistically distinct pathways, in which case interventions targeting IGF-1 signaling would fail to overcome the progeroid effects of mtDNA instability.

To distinguish between these possibilities, we reduced IGF-1 signaling in mice that carry an error-prone version of DNA Polymerase γ^18^ (*PolgA*^D257A^), the enzyme that replicates the mitochondrial genome. These mice accumulate mtDNA mutations at an accelerated pace^19,20^, which reduces their lifespan by 40% and leads to a wide variety of age-associated phenotypes, including sarcopenia, cardiomyopathy, anemia, and inflammation^18,21^. We crossed these mice into a background that is deficient for *Pappalysin 1* (*Pappa*), a metalloproteinase that increases IGF-1 bioavailability by cleaving IGF-binding protein 4 (IGFBP-4)^22^. In the absence of PAPPA, IGFBP-4 persists and sequesters IGF-1 away from receptors, thereby reducing IGF-1 signaling. When *Pappa* is deleted in WT mice, they exhibit a 35% increase in lifespan and a reduction in age-related pathologies^23–25^.

Surprisingly though, we found that *Pappa* deletion failed to extend the lifespan of *PolgA*^D257A^ mice, suggesting that the integrity of the mitochondrial genome is required for lifespan extension by reduced IGF-1 signaling.

Moreover, many of the pro-longevity programs normally initiated by *Pappa* deletion, including improved proteostasis, enhanced DNA repair and better telomere maintenance failed to initiate in *PolgA*^D257A^; *Pappa*^-/-^ mice. These observations suggest that mtDNA instability places a hard limit on mammalian lifespan, and that mitochondrial function is required for the successful activation of multiple longevity programs.

## RESULTS

### Lifespan extension by IGF-1 suppression depends on the integrity of the mitochondrial genome

To examine how IGF-1 signaling and mitochondrial mutagenesis interact to shape mammalian lifespan, we crossed *PolgA^D257A^* mice into a *Pappa^-/-^* background and compared the lifespan of WT mice to *PolgA*^D257A^ mice, *Pappa*^-/-^ mice, *PolgA*^D257A^; *Pappa*^-/-^ mice, *Pappa*^+/-^ mice, and *PolgA*^D257A^; *Pappa*^+/-^ mice. As expected, we found that *PolgA*^D257A^ mice exhibited a marked reduction in both median and maximum lifespan relative to WT controls, with all *PolgA^D257A^*mice expiring before they reached 17 months of age (**fig. 1A**), a timepoint when all WT mice were still alive. In contrast, the first *Pappa*^-/-^ mouse did not expire until it reached 28 months of age, consistent with their extended lifespan^12^. Surprisingly though, a complete deletion of the *Pappa* gene failed to extend the lifespan of *PolgA*^D257A^ mice (**fig. 1A**). Instead, *PolgA*^D257A^*; Pappa^-/-^* mice seemed to exhibit enhanced frailty, characterized by extensive weight loss as they aged (**fig. 1B-D,** for a weight distribution of young mice and their progression throughout their lifespan, see **fig. S1**). A heterozygous deletion of the *Pappa* gene failed to improve the lifespan of *PolgA*^D257A^ mice as well (**fig. 1A**), although this deletion was better tolerated, as indicated by their normalized body weight (**fig. 1B,C**). Taken together, these results indicate that the pro-longevity effects of IGF- 1 suppression are contingent upon the integrity of the mitochondrial genome.

**Figure 1.**
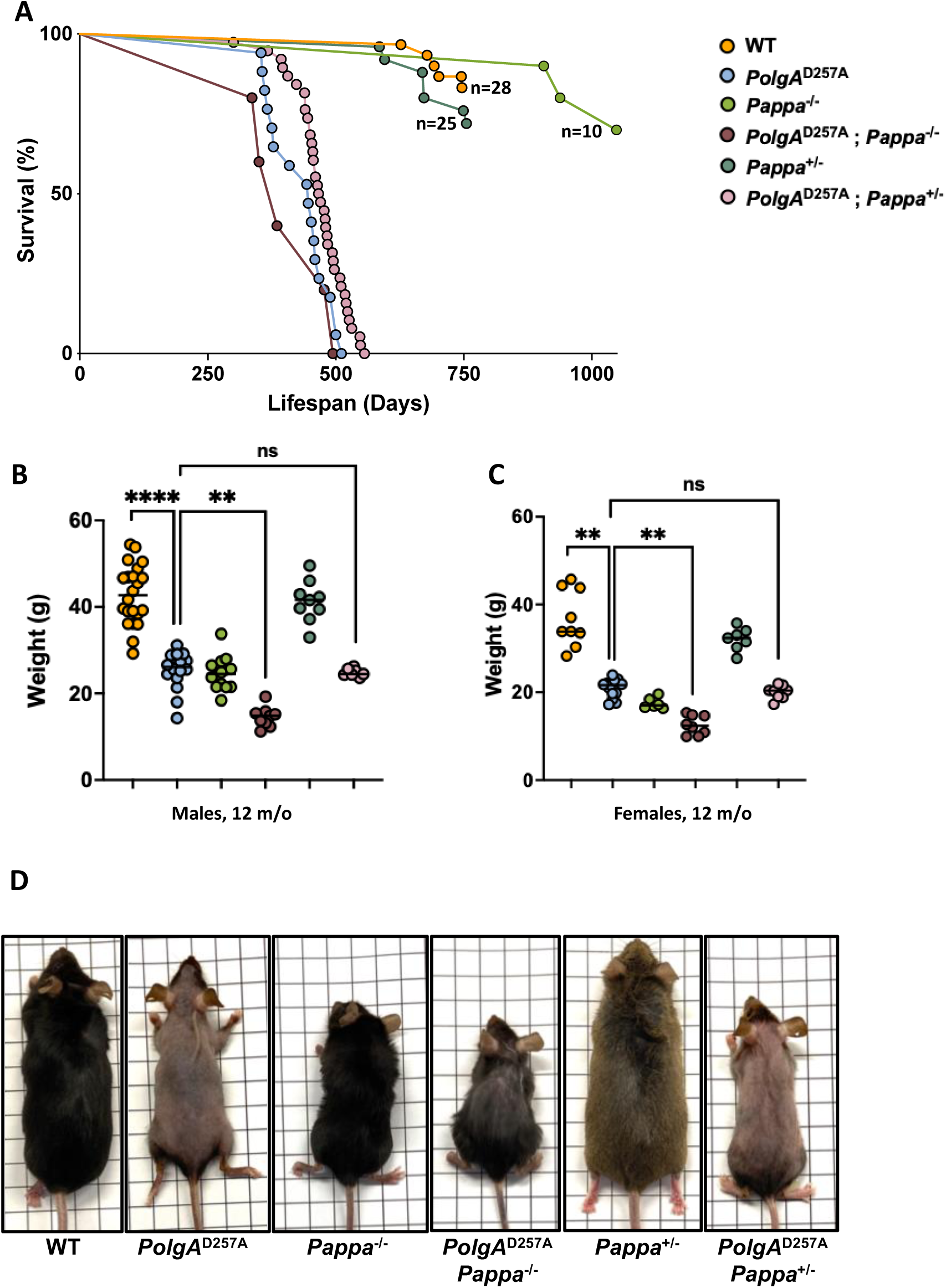
Lifespan, weight, size and appearance of WT, *PolgA*^D257A^ and *Pappa* mutant mice. **A.** The lifespan of *PolgA*^D257A^ is not rescued by deletion of the *Pappa* gene. **B-C**. A homozygous deletion of the *Pappa* gene leads to decreased weight and increased frailty in male and female *PolgA*^D257A^ mice (male n = 8-22/group, female n = 6-13/group). **D.** Appearance and size distribution of WT and mutant mice (grid size = 1 cm^2^).

### Deletion of *Pappa* rescues splenomegaly in *PolgA*^D257A^ mice

Next, we tested whether loss of *Pappa* could attenuate the age-related pathology of the mutator mice. Both male and female *PolgA*^D257A^ mice exhibit pronounced splenomegaly at 12 months of age (**fig. 2A-C**), and surprisingly, this phenotype was rescued in both sexes by homozygous deletion of the *Pappa* gene, even after we corrected for the reduced size of *Pappa*^-/-^ animals (**fig. 1D**). Interestingly, a heterozygous deletion of *Pappa* also rescued spleen size in male *PolgA*^D257A^ mice, although not in female mice, suggesting that male *PolgA*^D257A^ mice are more sensitive to IGF-1 depletion than females. These findings suggest that depletion of *Pappa* (loss of one copy of the *Pappa* gene) can partially recapitulate the impact of a full deletion (loss of both copies of the *Pappa* gene). These findings indicate that while *Pappa* deletion is insufficient to extend lifespan in *PolgA*^D257A^ mice, it *is* capable of rescuing distinct pathologies that are driven by mtDNA instability.

**Figure 2.**
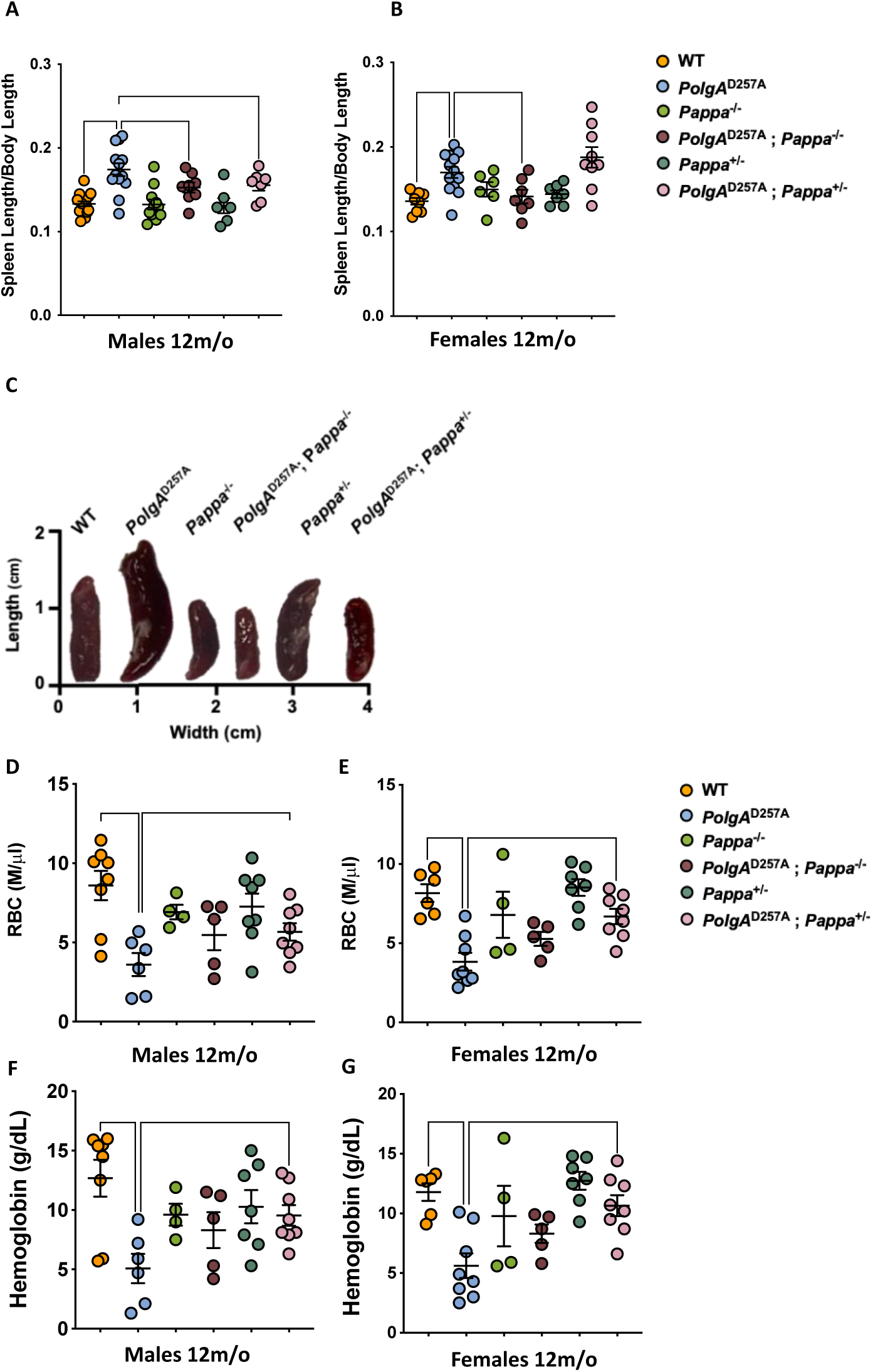
The splenomegaly and anemia of *PolgA*^D257A^ mice are partially rescued by deletion of *Pappa*. A-**C.** Male and female *PolgA*^D257A^ mice display splenomegaly. In both sexes, spleen size is reduced by deletion of two copies of the *Pappa* gene. In males, deletion of one copy of the *Pappa* gene is sufficient for spleen reduction as well. Spleen length was normalized to body length (male n = 7-17/group, female n = 6-13/group). **D-E.** Male and female *PolgA*^D257A^ mice exhibit reduced RBC counts, which is rescued in both sexes in *PolgA*^D257A^; *Pappa*^+/-^ mice. **F-G.** Male and female *PolgA*^D257A^ mice exhibit reduced hemoglobin content, which is rescued in both sexes in *PolgA*^D257A^; *Pappa*^+/-^ mice (male and female n = 4-8/group).

### Depletion of *Pappa* partially rescues anemia in *PolgA*^D257A^ mice

The reduced spleen size of *PolgA*^D257A^; *Pappa*^-/-^ mice prompted us to investigate whether other phenotypes associated with the *PolgA*^D257A^ allele were attenuated as well. A second characteristic phenotype of *PolgA*^D257A^ mice is anemia, which is thought to arise from multiple factors, including splenomegaly, which is an accelerant for the destruction of red blood cells (RBCs), impaired hematopoietic stem cell function, and increased damage to circulating erythrocytes^26^. Accordingly, we observed that old, but not young *PolgA*^D257A^ mice displayed a marked reduction in RBCs and hemoglobin content (**fig. 2D,E; fig. S2**). Excitingly, heterozygous deletion of the *Pappa* gene partially restored both the RBC levels and hemoglobin content in *PolgA*^D257A^ *Pappa^+/-^* mice (**fig. 2F,G**). In these experiments, as well as others (see below), *PolgA*^D257A^ *Pappa^+/-^* mice consistently outperformed *PolgA*^D257A^ *Pappa^-/-^* mice, suggesting a dose-dependent reduction in IGF-1 signaling is required for *PolgA*^D257A^ mice to achieve the greatest benefits.

### Depletion of *Pappa* partially rescues the sex-specific, inflammatory profile of *PolgA*^D257A^ mice

Chronic inflammation is a well-established consequence of mitochondrial dysfunction that is frequently driven by increased reactive oxygen species (ROS) production^27^, the release of damage-associated molecular patterns^28^, and activation of innate immune pathways such as the STING pathway^29,30^. In addition, inflammation is a major cause of anemia in patients with chronic diseases. Accordingly, we examined whether *PolgA*^D257A^ mice display elevated levels of inflammation as well and found that male *PolgA*^D257A^ mice exhibit significantly elevated serum levels of IL-2, IL-6, TNF-α, and IFN-γ, cytokines indicative of systemic inflammation and heightened splenic immune activity (**fig. 3**). Heterozygous deletion of Pappa significantly reduced IL-2 and IL-6 levels in these males, attenuating their inflammatory phenotype. Notably, *PolgA*^D257A^; *Pappa^+/-^* mice showed greater improvement than *PolgA^D257A^*; *Pappa^-/-^* mice, reinforcing the importance of IGF-1 dosage in ameliorating mtDNA disease. In addition, we noted that female *PolgA*^D257A^ mice displayed a muted inflammatory profile compared to male *PolgA*^D257A^ mice, indicating a sex-specific inflammatory response to mitochondrial mutagenesis. Most likely, this sexual dimorphism is due to the anti-inflammatory^31^ and anti-oxidant activity^32^ of estrogen, and other sex-linked factors^33^. Regardless, the limited inflammatory profile of female *PolgA*^D257A^ mice remained unchanged upon *Pappa* deletion (**fig. 3h**), highlighting the heightened sensitivity of male *PolgA*^D257A^ mice to IGF-1 modulation.

**Figure 3.**
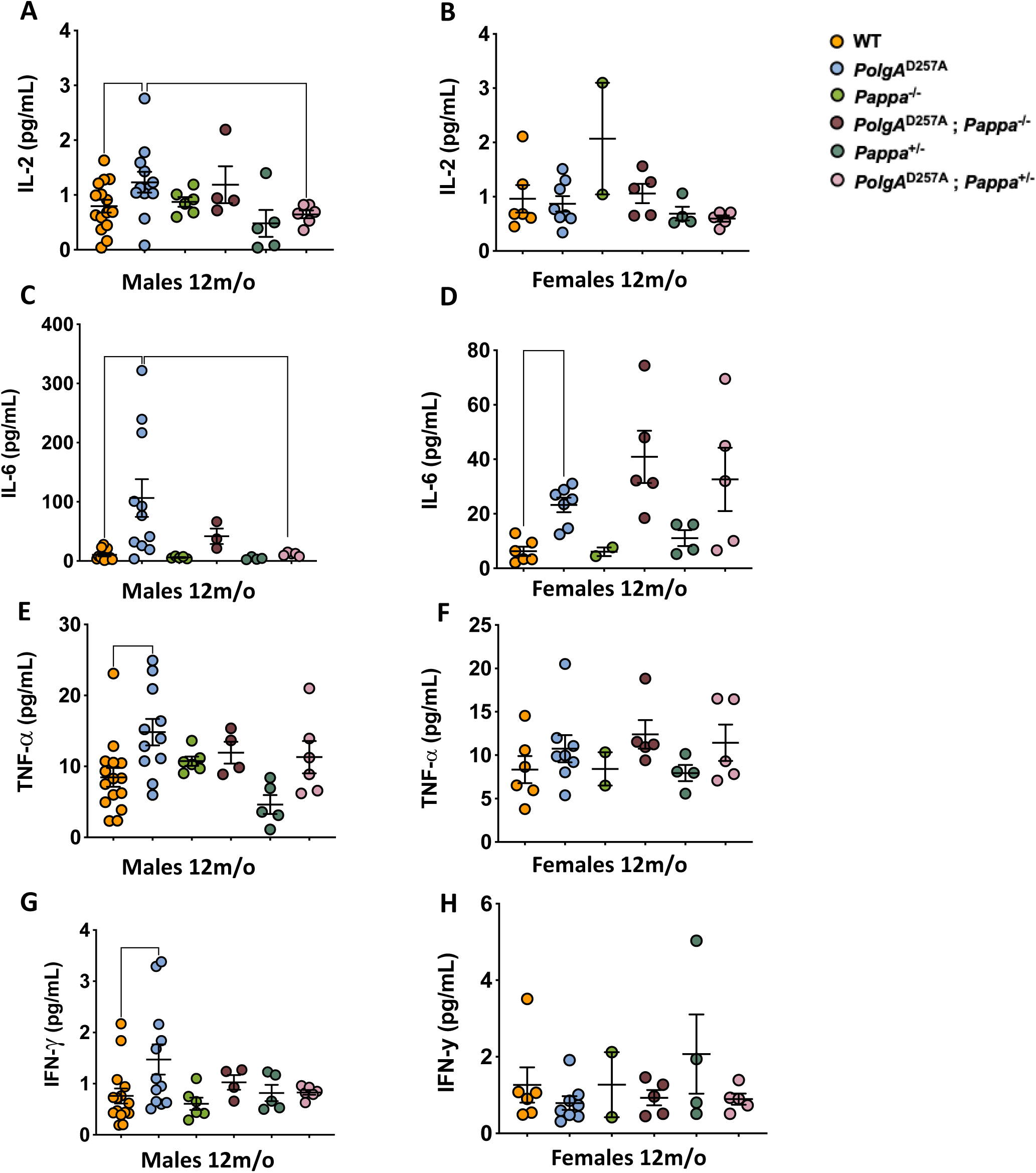
Inflammation in *PolgA*^D257A^ mice is partially rescued by depletion of *Pappa*. A, C, E, **G.** Male *PolgA*^D257A^ mice display increased levels of serum inflammation markers. IL-2 (**A**) and IL-6 (**C**) markers are rescued by deletion of one copy of the *Pappa* gene. **B, D, F, H.** Female *PolgA*^D257A^ mice only display increased levels of IL-6 among the evaluated cytokines, which is not rescued by *Pappa* deletion (**D**). (male n = 4-15/group, female n = 2-8/group).

### Depletion of *Pappa* improves muscle function in male *PolgA*^D257A^ mice

The primary consequence of mitochondrial mutagenesis is impaired energy production, which is particularly detrimental to energy-demanding tissues such as skeletal muscle^34^. As a result, aged (but not young, **fig. S3**) *PolgA^D257A^* mice exhibit significant reductions in both grip strength and treadmill endurance compared to WT type controls. These assays primarily assess type I (grip strength) and type II (endurance) muscle fiber performance, suggesting functional impairment across both fiber types. Consistent with our previous findings, we observed improved grip strength and endurance in *PolgA^D257A^; Pappa^+/-^* males, but not females and saw no improvement in *PolgA^D257A^; Pappa^-/-^* mice (**fig. 4A-D)**. Instead, *PolgA*^D257A^*; Pappa*^-/-^ mice tended to display reduced, rather than improved grip strength compared to *PolgA*^D257A^ mice, providing further evidence of their overall frailty.

**Figure 4.**
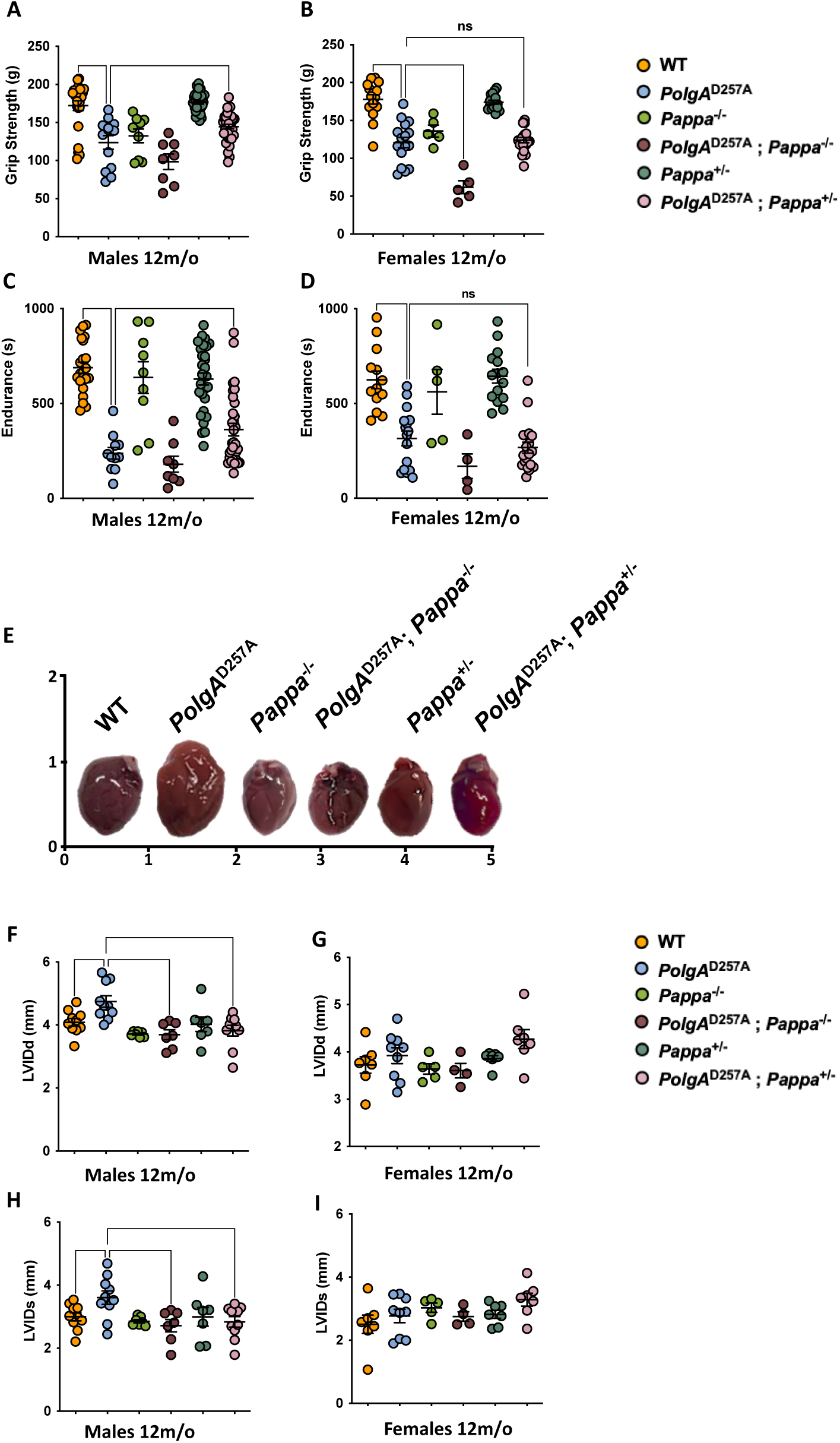
Impaired muscle and cardiac function in *PolgA*^D257A^ mice is partially rescued by deletion of *Pappa*. A-D. Male and female *PolgA*^D257A^ mice display reduced grip strength and endurance. In male mice, these physiological endpoints are improved upon deletion of one *Pappa* copy in male, but not female mice (male n = 8-32/group, female n = 5-19/group). **E-I.** Male, but not female, *PolgA*^D257A^ mice display increased LVID during diastole and systole. Heart size, LVIDd and LVIDs return to WT levels in male mice upon deletion of one or two copies of the *Pappa* gene (male n = 7-10/group, female n = 4-9/group).

### Deletion and depletion of *Pappa* improve the sex-specific cardiac dysfunction of *PolgA*^D257A^ mice

Like skeletal muscle, cardiomyocytes rely heavily on continuous energy production, making them particularly vulnerable to mitochondrial dysfunction^35,36^. In *PolgA*^D257A^ mice, extensive mitochondrial mutagenesis leads to significant cardiac hypertrophy and enlargement of the left ventricle—a phenotype that mirrors age-related cardiac pathology in humans. To determine whether *Pappa* deletion could mitigate these effects, we performed echocardiography on 12-month-old mice and found that male *PolgA*^D257A^ mice exhibited a significant increase in the heart size (**fig. 4E-I**), as well as the internal diameter of the left ventricle in both diastole (LVIDd) and systole (LVIDs), indicating structural remodeling and reduced cardiac efficiency. Similar to the inflammatory profile of the mice, we did not observe overt cardiomyopathy in female *PolgA*^D257A^ mice, suggesting a sex-specific impact of mtDNA instability on cardiac health, consistent with previous reports^37^. Because inflammation can drive cardiomyopathy^38^, it is possible that the reduced inflammation in female *PolgA*^D257A^ mice is mechanistically linked to their cardiomyopathy. We found that both *PolgA*^D257A^*; Pappa*^+/-^ and *PolgA*^D257A^*; Pappa*^-/-^ mice showed significant improvement in ventricular dimensions compared to *PolgA*^D257A^ males. These findings further confirm that depletion of *Pappa* partially phenocopies a complete deletion. Indeed, out of the 2,950 genes that are significantly altered in the hearts of male *Pappa*^-/-^ mice vs WT mice, 2,309 move in the same direction in *Pappa^+/-^*mice, albeit in a dose dependent manner (Spearman correlation 0.46, p=1.5x10^-154^, **fig. S4**). Together, these observations indicate that reduced IGF-1 signaling can partially restore cardiac function even in the context of a high mtDNA mutation burden

### The mutation frequency of mtDNA is not affected by deletion of the *Pappa* gene

To investigate the mechanism by which cardiac function is improved in male *PolgA^D257A^* mice following *Pappa* deletion, we tested whether reduced IGF-1 signaling lowers the mitochondrial mutation burden. Using duplex sequencing^39^, we generated high-resolution maps of the single base substitutions (**fig. 5A**), deletions (**fig. 5B**), and insertions (**fig. 5B**) that arose across the mitochondrial genome in 12 month old animals. However, the frequency of these mutation classes was unchanged among *PolgA*^D257A^, *PolgA*^D257A^*; Pappa*^+/-^, and *PolgA*^D257A^*; Pappa*^-/-^ mice, as was their mutation spectrum (**fig. 5C**) and the distribution of mutations across the mitochondrial genome (**fig. 5D,E**). Digital droplet PCR (**fig. 5F**) and qPCR (**fig. S5**) further showed that mtDNA copy number was unchanged among these genotypes. Thus, the improvements in cardiac function are not due to reduced mtDNA mutagenesis or altered mtDNA copy number, but must be mediated by processes that occur after mutation generation.

**Figure 5.**
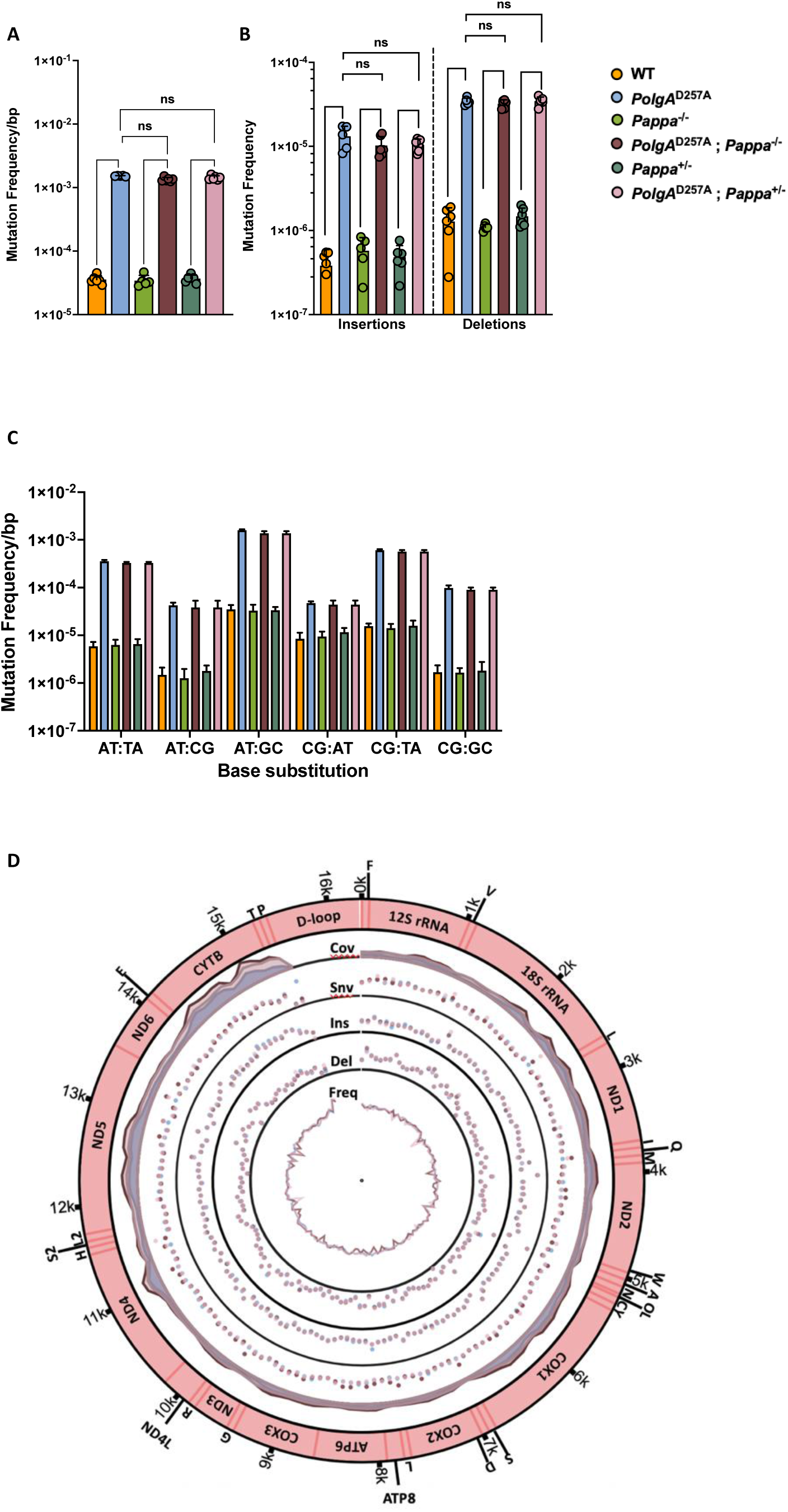
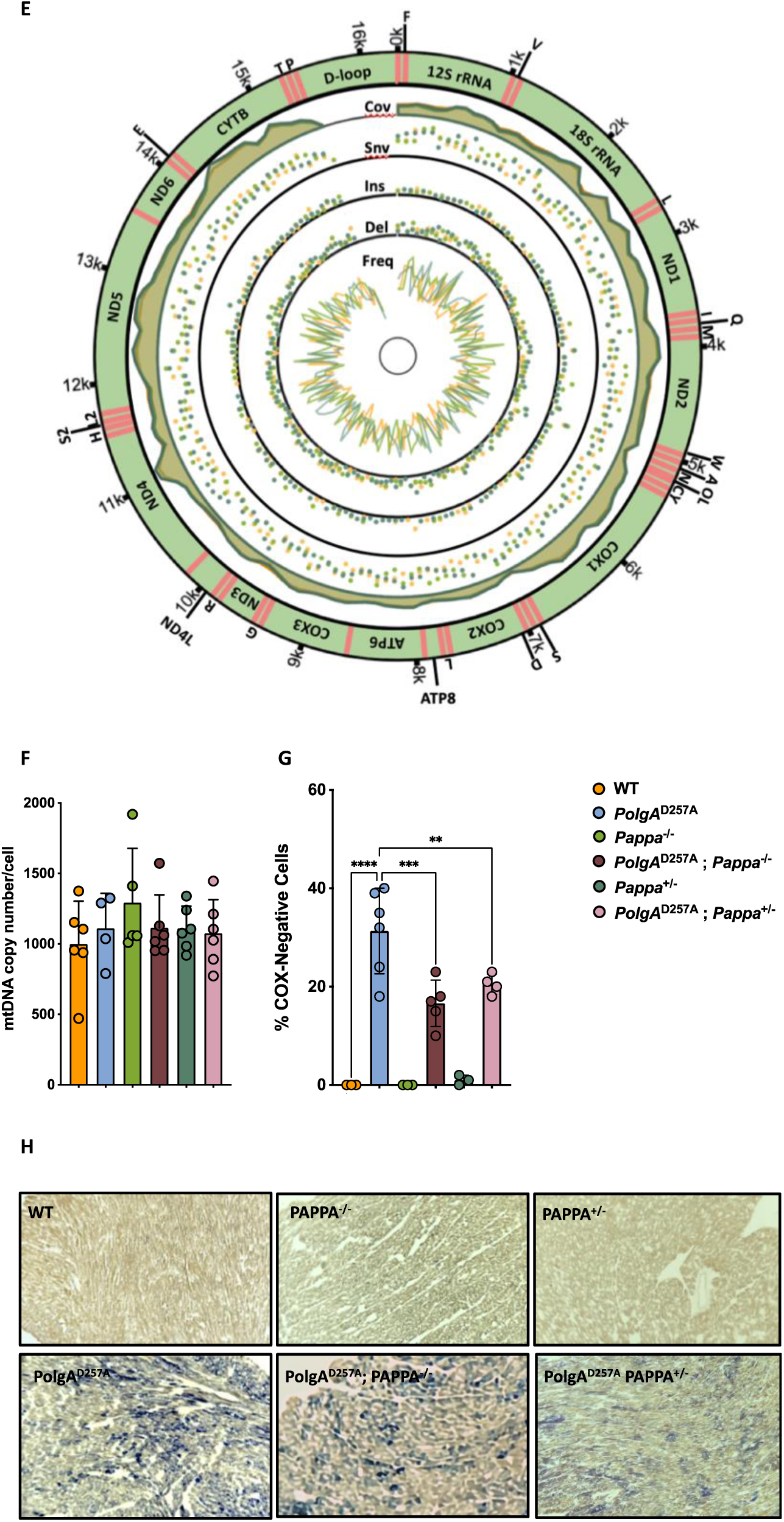
Mutagenesis in WT and mutant mice. A-C. Male *PolgA*^D257A^ mice, with or without *Pappa* deletion display identical mutation rates and spectra for all mutation classes. The same is observed for WT mice with or without *Pappa* deletion. **D,E.** Mutations were equally distributed across the mitochondrial genome (n = 5- 6/group). First track = coverage. Second track = single nucleotide variants. Third track = insertions. Fourth track = deletions. Fifth track = mutation frequency. **F.** mtDNA copy number was unchanged by deletion of the *Pappa* gene (n = 5-6/group). **G,H.** Deletion of the *Pappa* gene reduces the number of cardiomyocytes with clonally expanded mtDNA mutations. Blue cells depict cells without WT COX activity (n = 3/group).

### Deletion of *Pappa* slows clonal expansion of mtDNA mutations

While the mutation frequency remained unchanged upon *Pappa* deletion, the pathogenic consequences of these mutations do not manifest themselves until they clonally expand within individual cells. In both aging humans^34^ and *PolgA*^D257A^ mice^19^, these clonally expanded mutations disrupt cytochrome oxidase activity, impair ATP production and promote cardiomyopathy. To determine whether *Pappa* deletion influences this expansion process, we stained heart sections from 12-month-old mice for COX/SDH activity (**fig. 5H**), a technique that can identify cells in which mtDNA mutations have clonally expanded. While COX-negative cells were absent in WT, *Pappa^+/-^* and *Pappa^-/-^*hearts, *PolgA^D257A^* mice exhibited a substantial accumulation of COX-negative cells, consistent with widespread clonal expansion of mtDNA mutations. Remarkably, this phenotype was partially rescued in both *PolgA^D257A^; Pappa^+/-^* (p = 0.0046) and *PolgA^D257A^; Pappa^-/-^*(p = 0.0002) mice (**fig. 5G**). These findings suggest that although IGF-1 signaling does not affect the generation of mutations, it does affect the rate with which harmful mtDNA mutations clonally expand within cells, thereby preserving mitochondrial function and contributing to improved cardiac health in *PolgA^D257A^* mice.

### Transcriptomic profiling reveals molecular signatures of cardiomyopathy in male *PolgA*^D257A^ mice

To investigate the consequences of reduced homoplasmy in *PolgA*^D257A^; *Pappa*^-/-^ mice, we analyzed their heart tissue by RNA-seq. First, we established a baseline, transcriptomic profile of *PolgA*^D257A^ hearts (**fig. S6**) by comparing differentially expressed genes between *PolgA*^D257A^ and WT mice. This analysis confirmed that the hearts of *PolgA*^D257A^ mice are under substantial metabolic and inflammatory stress. Specifically, *PolgA*^D257A^ mice exhibited a 3-fold upregulation of *Gdf15* (p = 0.0005), a well-established marker of cardiac pathology, inflammation, and metabolic perturbation and a 7-fold increase in *Fgf21* (p = 0.03), which encodes a hormone that is released in response to metabolic stress. Consistent with a failing myocardium we also observed a 16- fold upregulation of *Myh7* (p < 0.0001) and a 3-fold upregulation of *Nppb* (p < 0.0001), two canonical markers of the "fetal gene expression program" that can be reactivated during cardiac dysfunction. These transcriptional changes were accompanied by a significant downregulation of genes that encode key components of the contractile apparatus, including *Myl1*, *Myl4*, *Myl7*, and *Mybphl*, all of which encode myosin light chains essential for sarcomere stability and contraction in striated muscle. Additionally, critical regulators of cardiac pacemaking and excitation-contraction coupling were downregulated, such as voltage-gated calcium (*Atp2b2, Cacna2d2*) and potassium channels involved in maintaining cardiac electrophysiology. Consistent with their inflammatory profile (**fig. 4**), we also observed a marked upregulation of chemokines that recruit innate immune cells, including *Ccl2* (3-fold), *Ccl8* (5-fold), *Cxcl5* (3-fold) and *Cxcl13* (>20-fold), potent attractants of monocytes, macrophages and neutrophils. Components of the complement system (e.g., *C4b*, 2.5-fold increase) and pattern-recognition receptors like *Tlr7* (2-fold) that are associated with the JAK-STAT signaling pathway were also elevated. Notably, a subset of these inflammatory genes was linked to the activation of the AIM2 inflammasome (enrichment p= 4×10⁻⁴), a complex that detects cytosolic double-stranded DNA (including mtDNA that leaks out of dysfunctional mitochondria) and drives innate immune activation^40^. Together, these findings establish a molecular signature of cardiomyopathy in *PolgA^D257A^* mice, characterized by metabolic stress, sarcomeric disorganization, electrophysiological instability, and robust activation of innate immune pathways.

### *Pappa* deletion partially normalizes the transcriptomic profile of male *PolgA*^D257A^ hearts

To assess the impact of *Pappa* deletion on the pathological transcriptomic landscape of *PolgA*^D257A^ mice, we then compared the changes in expression profile of *PolgA*^D257A^ vs. WT mice to *PolgA*^D257A^*; Pappa*^+/−^ vs. WT, and *PolgA*^D257A^*;Pappa*^−/−^ vs. WT animals. Remarkably, we found that deletion of one or two copies of the *Pappa* gene shifted the entire transcriptome of *PolgA*^D257A^ hearts closer to WT levels (**fig. 6A-C**). This normalization was most pronounced when we focused on the genes that are most dysregulated in *PolgA*^D257A^ animals. Out of the 252 genes that were significantly downregulated >2-fold in *PolgA*^D257A^ hearts, 217 moved closer to WT levels in *PolgA*^D257A^*; Pappa*^+/−^ mice (p < 0.0001) and 203 in *PolgA*^D257A^*; Pappa*^−/−^ mice (p < 0.0001, **fig. 6A**). A similar trend was observed among upregulated genes. Out of the 244 genes that were significantly upregulated >2-fold in *PolgA*^D257A^ hearts, 202 moved closer to WT levels in *PolgA*^D257A^*; Pappa*^+/−^ mice (p < 0.0001) and 167 in *PolgA*^D257A^*; Pappa*^−/−^ mice (p < 0.01, **fig. 6B**). In contrast, genes that were not dysregulated in *PolgA*^D257A^ mice showed no significant change when *Pappa* was deleted (**fig. 6C**).

**Figure 6.**
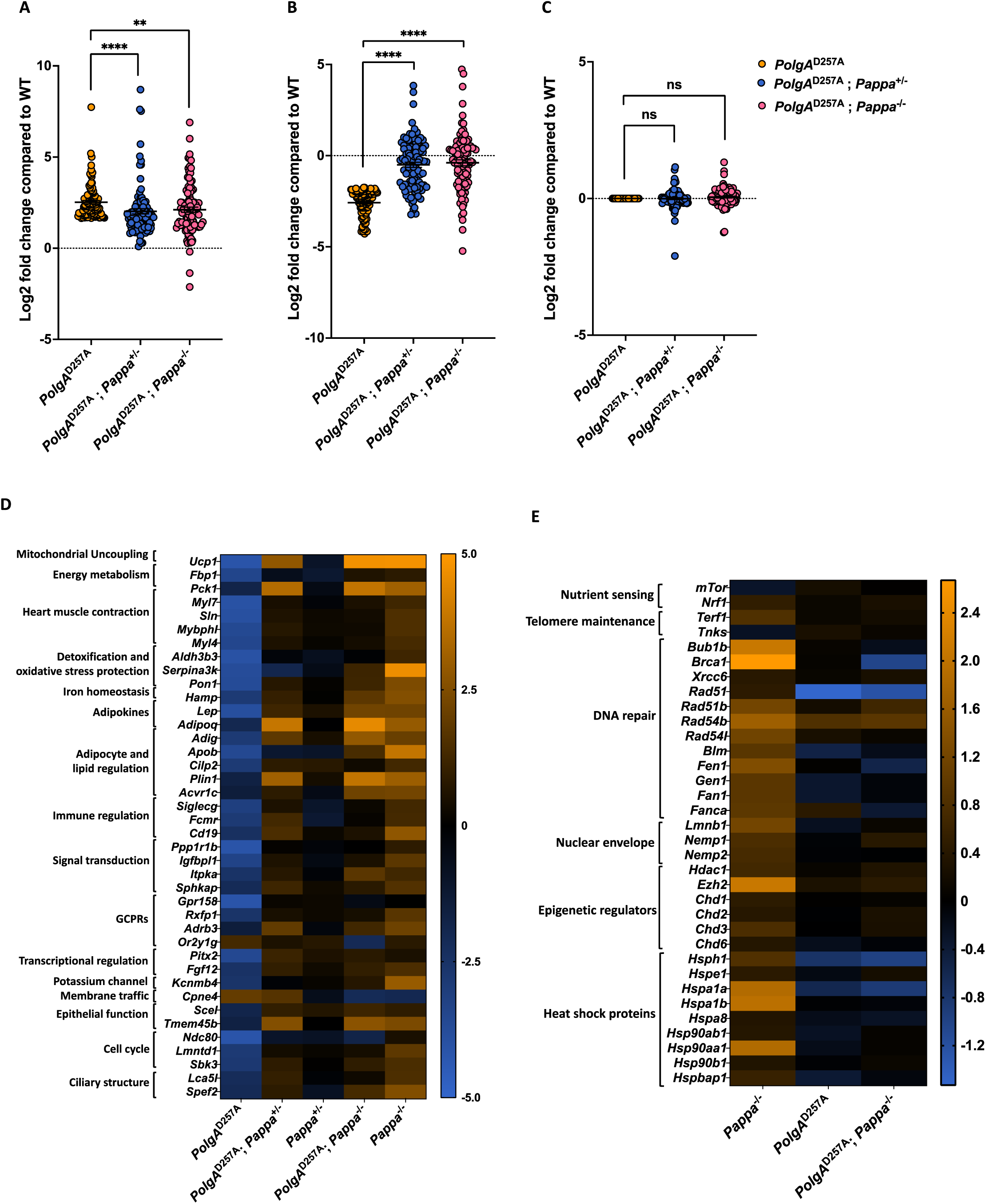
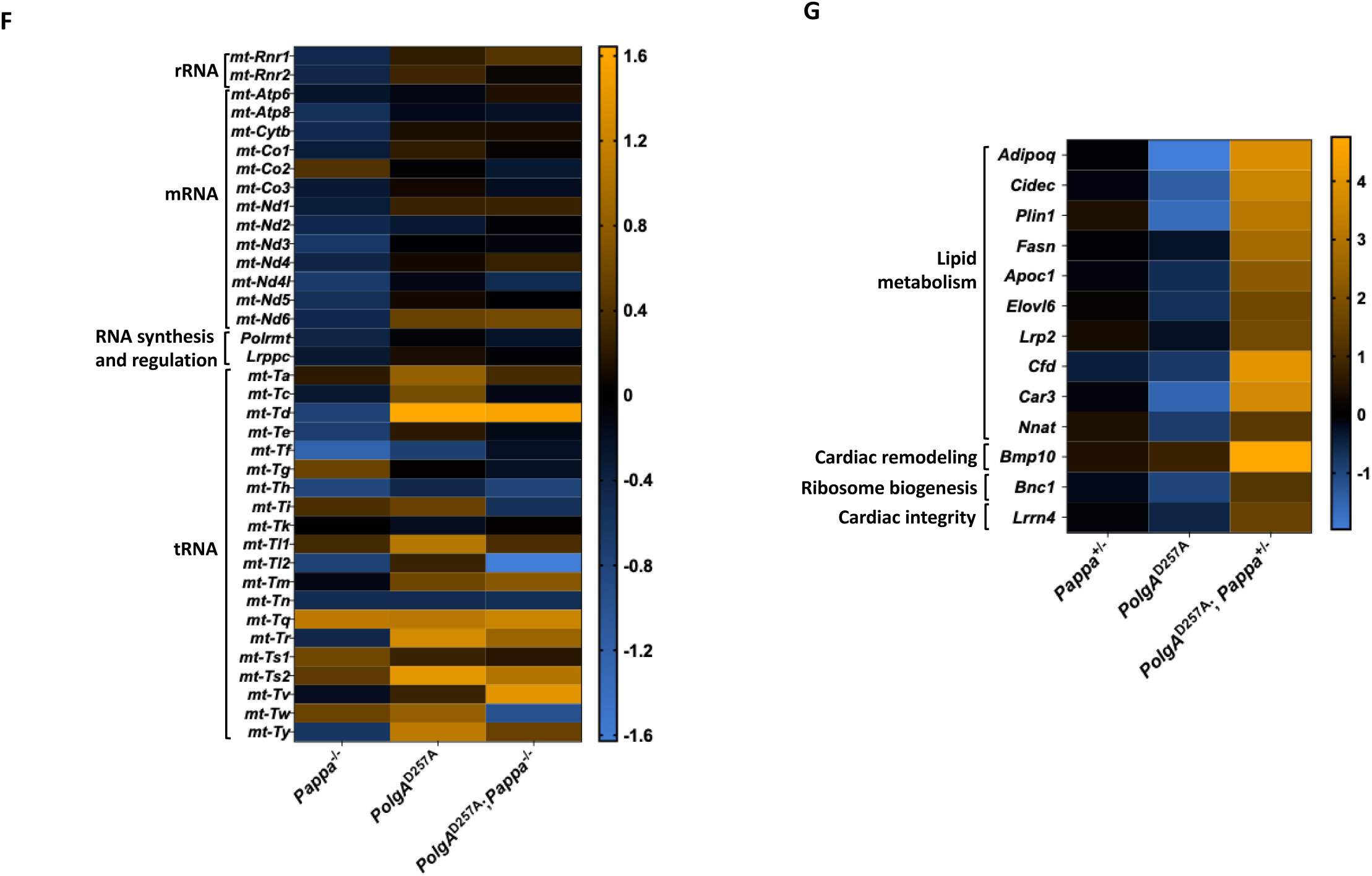
Transcriptomic analysis of WT and mutant mice. A-C. Loss of one or two copies of the *Pappa* gene normalizes the expression profile of *PolgA*^D257A^ mice. Genes that are upregulated (**A**) or downregulated (**B**) in *PolgA*^D257A^ mice are normalized upon deletion or depletion of *Pappa*. Genes that are unchanged in *PolgA*^D257A^ mice remain unchanged after complete or partial loss of *Pappa* (**C**). **D.** Gene expression patterns suggest that impaired cardiac and metabolic function are rescued by deletion and depletion of the *Pappa* gene, drastically shifting the expression patterns away from the dysregulation observed in *PolgA*^D257A^ mice. Interestingly, loss of one copy of *Pappa* in *PolgA*^D257A^ mice changes the transcriptional profile in the direction of *Pappa*^-/-^ compared to *Pappa^+/-^*, suggestive of implementation of pathways to rescue the negative consequences of mtDNA mutagenesis. **E.** Pro-aging related gene programs are dampened by deletion and depletion of *Pappa*. **F.** Complete loss of *Pappa* does not restore expression patterns of mitochondrial transcripts in the *PolgA*^D257A^ background. **G.** Lipid metabolism related genes are not upregulated in *Pappa*^+/-^ mice or *PolgA*^D257A^ mice, but are upregulated in *PolgA*^D257A^; *Pappa^+/-^* mice, suggesting context dependent expression patterns (n = 6/group).

### *Pappa* deletion restores metabolic flexibility and cardiac function in male *PolgA*^D257A^ hearts

To identify the pathways most responsible for the improved cardiac health of PolgA^D257A^ mice, we filtered the transcriptome for genes that change ≥8-fold in expression upon deletion of either one, or two copies of the *Pappa* gene. This analysis showed that cardiac health was improved at multiple levels of scale. At a functional level, we found that key cardiac contractile genes (*Myl7*, *Myl4*, *Mybphl*, *Sln*) that were strongly downregulated in *PolgA*^D257A^ mice, were robustly restored by *Pappa* deletion, suggesting a rescue of cardiac contractile function (**fig. 6D**). This functional rescue was accompanied by widespread metabolic reprogramming. Although heart tissue primarily relies on fatty acid oxidation^41^, it can switch to glucose utilization under stress^42^. Two genes that enable this metabolic flexibility are PCK1 the rate limiting enzyme for gluconeogenesis, and FBP1^43^, which are significantly downregulated in *PolgA*^D257A^ hearts, indicating metabolic rigidity. Deletion of *Pappa* reactivates these genes, indicating enhanced gluconeogenesis and glucose mobilization, which improves the metabolic flexibility of the heart. Systemically, *PolgA*^D257A^ mice exhibit reduced expression of the adipokines leptin (*Lep*) and adiponectin (*Adipoq*), consistent with global metabolic dysfunction. *Pappa* deletion reversed this trend, restoring expression of both hormones and correcting this endocrine imbalance. Restoration of *Adipoq* is particularly noteworthy, given its role in insulin sensitivity, mitochondrial biogenesis, and antioxidant defenses^44^, factors that influence organismal resilience at multiple levels of scale. Further transcriptomic shifts supported improved lipid handling. *Pappa* deletion upregulated Adipogenin (*Adig*), which promotes adipocyte differentiation^45^, and *Plin1*, which stabilizes lipid droplets and prevents lipotoxicity^46^. These changes occurred alongside increased expression of *Adrb3*^47^ and *Acvr1c*^48^, regulators of lipid mobilization and adipocyte metabolism. Notably, *Adrb3* was co-rescued with *Ucp1*, suggesting improved thermogenesis and mitochondrial uncoupling and the activation of stress-protective metabolic programs. Together, these changes suggest that *Pappa* deletion enables *PolgA*^D257A^ hearts to shift from metabolic exhaustion towards a more flexible, energy and stress-resilient metabolic state.

### The *PolgA*^D257A^ allele blocks anti-aging programs triggered by *Pappa* deletion

To understand why *Pappa* deletion can improve cardiac function, but not extend lifespan in *PolgA*^D257A^ mice, we analyzed the expression of genes that are involved in key anti-aging pathways (**fig. 6E**). One hallmark of lifespan extension is reduced mTOR signaling^49^. Accordingly, *Pappa*^-/-^ mice show reduced *mTor* expression (p= 0.046), while short-lived PolgA^D257A^ mice exhibit increased expression (p= 0.01) ; however, *mTor* reduction was partially blocked when *Pappa* was deleted in *PolgA*^D257A^ mice, limiting its impact on longevity. Another hallmark of longevity is improved telomere maintenance^1^. Consistent with their extended lifespan, *Pappa*^-/-^ mice, show upregulation of *Terf1* (p=0.0004) and downregulation of tankyrase (p=0.036), two changes that shield telomeres from damage and recombination and prevent them from triggering cell cycle arrest^50^. Importantly, neither gene changed expression when *Pappa* was deleted in a *PolgA*^D257A^ background. In addition, *Pappa*^-/-^ mice exhibit overexpression of multiple DNA repair genes that are involved in double strand break repair and telomere maintenance, a key feature of longevity programs^51^. However, these genes were not overexpressed in *PolgA*^D257A^; *Pappa*^−/−^ mice, demonstrating that the *PolgA*^D257A^ allele blunts, or blocks this longevity interventions as well. Another feature of long-lived animals is their ability to maintain their epigenetic landscape, which is normally eroded by reduced expression of epigenetic regulators like *Hdac1*. Accordingly, *Pappa*^-/-^ mice overexpress multiple chromatin regulators, including *Hdac1* (p=0.005), *Ezh2* (0.0002), *Chd1* (p=0.03), *Chd2* (p=0.02), *Chd3* (p=0.0007) and *Chd6* (p=0.049), none of which were overexpressed in *PolgA*^D257A^; *Pappa*^−/−^ mice. As cells age, the structure of their nuclear envelope tends to deteriorate, a change that is frequently attributed to reduced expression of Lamin B (*Lmnb1*)^52,53^. Consistent with their increased longevity, *Pappa*^-/-^ mice overexpress *Lmnb1* (p=0.002) together with *Nemp1* (p=0.006) and *2* (p=0.042), two structural proteins of the nuclear envelope; however, we found no overexpression of these genes in *PolgA*^D257A^; *Pappa*^−/−^ mice. As part of a shift from anabolic to catabolic metabolism, deletion of *Pappa* decreases expression of protein coding genes from the mitochondrial genome, which may reduce mitochondrial dysfunction and ROS production (**fig. 6F**). This reduction was mediated by reduced expression of the mitochondrial RNA polymerase (*Polrmt*, p=0.03) and LRPPC (p=0.06), a protein that selectively protects mitochondrial mRNAs, but not tRNAs^54,55^. Accordingly, the expression of tRNAs was not systemically reduced in *Pappa*^-/-^ mice. None of these changes were observed in *PolgA*^D257A^; *Pappa*^−/−^ mice. Another hallmark of long-lived cells is improved proteostasis^1^. Again though, we found that although *Pappa* deletion significantly upregulated numerous primary heat shock proteins, these genes were not upregulated in *PolgA*^D257A^; *Pappa*^-/-^ mice. Indeed, we found that out of the 32 primary heat shock proteins (HSPs) we detected, 27 were downregulated in *PolgA*^D257A^ mice (**fig. S7A**) and none of these genes were rescued in *PolgA*^D257A^; *Pappa*^−/−^ mice. Notably, this pattern was not observed in the DnaJ-family HSP40 of co-chaperones (**fig. S7B**), which, unlike the primary chaperones, are ATP-independent^56^. Thus, loss of mtDNA integrity, and reduced ATP production specifically inhibit the expression of ATP-dependent chaperones, contributing to proteostasis failure.

### *Pappa* deletion triggers context-dependent metabolic rewiring

Aging cells frequently exhibit not one, but multiple hallmarks of the aging process. These combinations could lead to complex, context-dependent interactions that result in novel gene expression states that neither hallmark produces alone. To identify such interactions between mitochondrial mutagenesis and IGF-1 signaling, we filtered the transcriptome for genes that remain unchanged upon deletion of *Pappa* in WT mice (*Pappa*^+/-^ mice), but were strongly altered (>4-fold) in *PolgA*^D257A^; *Pappa*^+/-^ mice. Of the 13 genes that were identified by this method, 7 were directly involved in lipid metabolism, storage, or mobilization (**fig. 6G**). These included *Adipoq*, a protective adipokine in metabolic disease and inflammation; *Cidec* and *Plin1*, which regulate lipid droplet formation and adipocyte energy storage^46,57^, and *Apoc1*, a lipid- and immune-regulatory apolipoprotein^58^. Finally, we identified *Fasn*, an enzyme that is central to fatty acid synthesis^59^; *Elovl6*, which catalyzes the elongation of long-chain fatty acids^60^, and *Lrp2*, a multifunctional endocytic receptor that internalizes lipoproteins, insulin, leptin, and amyloid-beta^61^. These results reveal a striking lipid-centric response to a partial deletion of *Pappa* that only emerges in the presence of mitochondrial dysfunction, underscoring how aging hallmarks can interact with each other to produce context-dependent transcriptional programs. Notably, some of these interactions also occur when both copies of *Pappa* are deleted, demonstrating that mitochondrial mutations can sensitize cells to modulation by *Pappa* deletion and offering a plausible explanation for why partial *Pappa* deletion can improve the phenotype of *PolgA*^D257A^ mice as well. A similar effect is seen when column 2 and 3 of **figure 6D** are compared.

## DISCUSSION

Aging is a multifactorial process, characterized by the simultaneous emergence of multiple, interconnected molecular and cellular hallmarks. In naturally aging organisms, these hallmarks do arise independently, but instead accumulate and interact within the same tissues and cells over time. As a result, there is a growing recognition that the hallmarks of aging are critically intertwined, and that to fully understand the aging process, it will be essential to tease apart these complex interactions.

In this study, we investigated how mtDNA instability interacts with IGF-1 signaling to shape mammalian lifespan and healthspan. Suppression of IGF-1 signaling through *Pappa* deletion extends lifespan in WT mice by triggering well-known pro-longevity programs, including enhanced stress resistance, DNA repair, proteostasis, and metabolic reprogramming. However, the same intervention fails to extend lifespan in *PolgA*^D257A^ mice, which accumulate mtDNA mutations at an accelerated pace. This observation echoes prior studies that showed that calorie restriction^62^, exercise^63–66^, or overexpression of a mitochondrially targeted catalase^67^, did not extend the lifespan of *PolgA*^D257A^ mice either. Taken together, these studies support that a hierarchy exists among the hallmarks of aging, where mtDNA integrity overrides the impact of longevity interventions, such as IGF-1 signaling, nutrient sensing, exercise and oxidative stress to impose a hard limit on mammalian lifespan.

Although *Pappa* deletion did not extend the lifespan of *PolgA*^D257A^ mice, it did attenuate a wide range of age-related phenotypes, including splenomegaly, anemia, chronic inflammation, skeletal muscle decline, and cardiomyopathy. These findings echo those with a mitochondrially targeted catalase, which failed to extend lifespan as well, but improved the cardiomyopathy in *PolgA*^D257A^ mice^35,67^. The benefits we observed were especially pronounced in male mice, which seemed to suffer more from mtDNA instability than female mice, possibly due to the anti-oxidant and anti-inflammatory effects of estrogen^68^. Surprisingly, these improvements were most evident in mice with partial *Pappa* deletion. This dose-dependent effect suggests that complete loss of *Pappa* may excessively suppress IGF-1 signaling in the context of high mitochondrial stress, where some degree of anabolic signaling remains necessary to maintain cellular homeostasis. Our transcriptomic analyses demonstrate that the health improvements in *PolgA*^D257A^; *Pappa*^+/-^ mice were mediated, at least in part, by context-dependent interactions between mtDNA instability and IGF-1 signaling. In other words, we observed gene expression states in *PolgA*^D257A^; *Pappa*^+/-^ mice that were not observed in either genotype alone. Such context-dependent effects underscore how important the interactions between the hallmarks of aging are: the outcome is not merely the sum of two interventions, but a new state shaped by their interplay.

This principle was most clearly illustrated by the failure of *Pappa* deletion to activate canonical longevity pathways in *PolgA*^D257A^ mice. In WT mice, *Pappa* deletion upregulates a suite of anti-aging pathways that improve DNA repair, telomere maintenance, proteostasis and chromatin integrity. However, in the presence of mtDNA mutations, most of these pathways are either blocked or blunted, demonstrating that the integrity of the mitochondrial genome is pre-requisite for lifespan extension by reduced IGF-1 signaling. Similarly, previous research has shown that proteostasis^69^, DNA repair^70^ and other molecular mechanisms are pre-requisites for lifespan extension by reduced IGF-1 signaling as well. Our observations now suggest that within this framework, mtDNA integrity is not simply one of the many hallmarks of aging, but rather the foundation upon which others are built. And when that platform is broken, downstream hallmarks like proteostasis or DNA repair cannot be engaged by typical means, no matter how favorable the upstream signaling may be.

This insight has major implications for the development of anti-aging therapies. It suggests that interventions that target nutrient-sensing pathways may fail—or even backfire—when applied to organisms or tissues with high levels of mitochondrial damage. As such, the next generation of geroprotective treatments must be tested in diverse models of aging, including those that combine multiple hallmarks, to better understand the scope and boundaries of their efficacy.

Because mtDNA mutations impose a ceiling on mammalian lifespan, it is vital that these treatments also address the stability of the mitochondrial genome. While a reduction in IGF-1 signaling did not alter the frequency of mutations in WT or *PolgA*^D257A^ mice, it did slow the pace with which they reached homoplasmy. Thus, although it may not be possible today to reduce mitochondrial mutagenesis in human cells, our data shows that it may already be possible to curtail the impact of mtDNA mutations on mammalian healthspan by slowing their clonal expansion.

While the precise mechanism by which *Pappa* influences clonal expansion of mtDNA mutations remains uncertain, several plausible explanations can be proposed. First, the progression of mtDNA mutations toward homoplasmy is likely influenced by mitochondrial genome replication and segregation dynamics, two processes that can accelerate genetic drift, especially during cell division when mutant genomes may be unequally partitioned between daughter cells. Because *Pappa* deletion suppresses growth-related pathways these changes may reduce mitochondrial workload and mtDNA replication frequency, thereby slowing genetic drift in post-mitotic cells and limiting mitochondrial segregation via reduced cell turnover. Second, *Pappa* deletion enhances a wide variety of maintenance processes, which could lead to improved autophagic and mitochondrial quality control mechanisms that selectively eliminates dysfunctional mitochondria and curbs the expansion of mutant clones. Third, emerging evidence suggests that damaged mtDNA is often degraded rather than repaired^71^. In that context, the enhanced antioxidant capacity observed in *Pappa*-deficient *PolgA*^D257A^ mice may lessen oxidative mtDNA damage, decreasing the need for mtDNA turnover and potentially stabilizing the mitochondrial genome by limiting replication-driven amplification of existing mutations.

Regardless, these findings provide a compelling example of how the interplay between distinct hallmarks of the aging process can fundamentally alter the outcome of otherwise beneficial interventions. They reveal that the efficacy of anti-aging strategies like IGF-1 suppression is not absolute, but context-dependent. They are contingent on the integrity of underlying systems, including proteostasis and DNA repair. Without an intact mitochondrial genome though, these pathways cannot be engaged, indicating that mtDNA integrity is upstream from these critical anti-aging pathways. More broadly, our results underscore the need for a more integrated model of aging, one that considers not just individual pathways but their interactions, hierarchies, and points of failure. By mapping these interactions, we can better anticipate the limitations of existing interventions and design next-generation therapies that are robust to the complex biology of aged tissues. In this light, strategies that target the expansion of mtDNA mutations—rather than their origin—may offer a powerful new axis for preserving tissue function and extending healthspan, even when the underlying genomic damage cannot be undone.

## METHODS

### Transgenic Mice

*PolgA*^D257A^ mice were purchased from JAX (strain #:017341), while *Pappa*^-/-^ mice were a generous gift from Dr. Cheryl Conover^12^. Male *PolgA*^D257A/+^; *Pappa*^+/-^ and female *PolgA*^D257A/+^; *Pappa*^+/-^ were bred to generate mice the genotypes described in this study. To limit mtDNA mutations from being inherited through the germline, separate *PolgA*^D257A/+^; *Pappa*^+/-^ x WT C57Bl6/j crosses were set up to generate “first generation” *PolgA*^D257A/+^; *Pappa*^+/-^ females to be used for breeding. At 21 days, mice were weaned, ear-punched and two PCR reactions were run to genotype each mouse. The first reaction genotyped the *PolgA*^D257A^ allele, using primers F: 5’- GCCTCGCTTTCTCCGTGACT-3’ and R: 5’-GGATGTGGCCCAGGCTGTAACTCA-3’. To genotype the *Pappa* allele, primers F_common_: 5’- TAAGCAGGGGTGGGTCCTTT-3’, F_neo_: 5’- TCGCCTTCTATCGCCTTCTTG-3’, and R: 5’- CACTCCTCAGCTTCGGCTTTCA-3’ were used. PCR reactions were run on agarose gels by gel electrophoresis and imaged before analysis. Mice were assayed at 3 months or 12 months of age for experimental studies, while a subset of mice were aged for lifespan experiments and weekly body weight measurements. Animals were scored by our lab and USC’s Department of Animal Resources veterinarians to establish humane endpoints. Mice were euthanized by CO_2_ exposure.

### Grip Strength

Grip strength was assessed using a horizontally mounted bar attached to a sensor (TSE-Systems, 303500- M/E1) designed to measure forelimb strength. Each mouse was allowed to securely grasp the bar before being swiftly and steadily pulled backward. Measurements were recorded only if the mouse let go with both forelimbs at the same time. Ten measurements were taken per animal and averaged to obtain a final value.

### Endurance

Mice were acclimated to the treadmill instrument (TSE 303401-M-04/C) 24 hours before testing. During acclimation, animals were placed on the stationary treadmill for five minutes, followed by an increase in speed to 2m/min for five additional minutes. Mice were then tested once each day on the two days following acclimation. The treadmill protocol consisted of 1 m/min for 1 minute, followed by a 1m/minute increase every minute until exhaustion. Time of exhaustion was recorded in seconds. Mice that refused to run would feel a mild shock at the back of the treadmill. The values recorded for each mice over both test days was averaged and reported.

### Echocardiography

One week prior to dissection, mice were transported to the molecular imaging center at USC for echocardiography using the VisualSonics Vevo 3100, MX 550 transducer 22-55 MHz. Mice were anesthetized with 2% isoflurane and a depilatory cream (Nair) was used to remove fur prior to echocardiography. Mice were placed in the supine position onto the warmed platform to maintain optimal physiological conditions and their limbs were taped onto the metal EKG leads. Heart rates were monitored and generally maintained at 400–500 beats per minute. Warmed echocardiography gel was placed on the shaved chest and the heart was imaged with a 30 MHz transducer. By placing the transducer along the long-axis of LV and directing it to the right side of the neck of the mouse, two-dimensional LV long-axis can be obtained. The transducer was then rotated clockwise by 90°, and the LV short-axis view was obtained. Transmitral inflow Doppler spectra were recorded in an apical 4-chamber view by placing the sample volume at the tip of the mitral valves. After the scans were concluded, the residual gel was removed, and the mouse was returned to the cage for recovery. Images were subsequently analyzed using the VevoLab software.

### Tissue Collection and Analysis

Upon euthanasia, mice were measured, photographed, and the heart was punctured for blood draws for blood composition experiments and serum isolation. Tissues were dissected, placed on a grid sheet for measurements and photography and cut into 4 pieces. One piece was stored in formalin, one piece was frozen in Optimal Cutting Temperature (OCT) compound and two pieces were flash frozen in liquid nitrogen for DNA and RNA extractions. Formalin-fixed tissue was left at room temperature overnight before being washed twice with PBS and stored at 4°C in 70% EtOH. Serum was collected by letting blood clot for 30 min, followed by a 10 minute spin at 1,000g at 4°C. Serum was then collected by aspirating the supernatant and stored at -80°C. IFN-γ, IL2, IL6 and TNF-α levels in serum were measured with commercial immunoassays using a custom V-PLEX Mouse Biomarker Kit (Meso Scale Discovery, Rockville, MD). Blood composition was measured using 40 μl of blood collected in EDTA tubes using the Hemavet 950FS instrument.

### Duplex-Sequencing

DNA was extracted from the hearts of experimental mice (n= 5-6/group) using the Qiagen DNeasy Blood and Tissue Kit (cat# 69506), quantified by Qubit 1x dsDNA hs assay kit, and stored at -20C. 500ng of DNA per sample was fragmented using the Biorupter Pico (30 seconds ON, 90 seconds OFF, 6 cycles) and size was measured using the Agilent 4200 TapeStation instrument (fragment size ∼300bp). Published protocols^39,72^ were followed with minor adjustments. Briefly, DNA was end-repaired and adapters (a generous gift from the Kennedy Lab) were ligated to DNA fragments using the Ultra II DNA Library Preparation Kit from NEB (cat# E7103) samples were cleaned up and libraries were quantified by qPCR using SYBR Green iTaq Supermix (Biorad cat#: 1725124) compared to previously generated libraries with target family sizes and on-target efficiencies. This quantification enables us to input the correct number of molecules into the pre-enrichment PCR depending on mtDNA content for each sample to sequence at the target parameters (depth, target raw reads, family size, on-target reads). Libraries were then enriched for mtDNA sequences using the xGen Hybridization Capture Assay (IDT) with a custom Discovery Pool of biotinylated probes (**Suppl. File 1**), along with their protocol. Final libraries were quantified, pooled and sequenced on a NovaSeq 6000 S2 kit (150PE). The standard duplex-seq pipeline was used to analyze the sequencing data, https://github.com/Kennedy-Lab-UW/Duplex-Seq-Pipeline. Mutation data are attached as supplementary files (**Suppl. File 2**).

### mtDNA Copy Number Measurements

DNA extracted for duplex-sequencing experiments was used for copy number determination. By quantitative PCR, relative mtDNA copy number was measured using 50ng of input DNA and primers targeting mitochondrial ND1 (F: 5’-GCCTGACCCATAGCCATAAT-3’ and R: 5’-TATTCTACGTTAAACCCTGA-3’) and nuclear RPP30 (F: 5’-GCAACCGGAACATAGAGACA-3’ and R: 5’-CTGGCCTTGGAATGGGTAAT-3’). iTaq Universal SYBR Green Supermix was used in these reactions and the protocol was followed accordingly (Biorad cat#: 1725124). By droplet digital PCR (ddPCR), BioRad ddPCR copy number assay primers/probe for Rpp30 (HEX, Assay ID: dMmuCNS822293939) and for ND1 (FAM, Assay ID: dMmuCNS343824284) were used with 2x ddPCR Supermix for Probes (no dUTP) (Biorad cat#: 186-3023).

### RNA-seq

Heart tissue from 12 month old, male mice (n = 6/group) was dounced in Trizol/Chloroform and homogenized with with zirconia beads for 20 minutes at 4°C. Samples were then at 10,000g for 5 minutes and supernatant purified according to the RiboPure Yeast RNA Purification Kit (Invitrogen, cat#: AM1926). RNA was DNase treated and mRNA was enriched using the GenElute Direct mRNA miniprep kit (Sigma, cat#: DMN70-1KT) and RNA-seq libraries were generated using the SMARTer Stranded RNA-Seq Kit (Takara Bio, cat#: 634838) and sequenced on a NovaSeq 6000 SP PE150 kit. Sequencing reads were trimmed using fastp^73^ (version 0.23.2) with the following parameters: "--trim_poly_g --poly_g_min_len 6 --length_required 120". Trimmed reads were then used to obtain transcripts abundance with Kallisto quant^74^ (version 0.46) against the Ensembl r96 annotation for Mus musculus. Abundance estimates from Kallisto were then analyzed in R with the package DESeq2^75^ following the standard pipeline to identify genes differentially expressed relative to the WT with FDR < 0.05 (correction for multiple testing). Sequencing data were deposited to SRA under accession PRJNA1271490.

### COX-SDH Staining

Frozen OCT blocks containing heart tissue were sectioned onto slides using a Leica CM 1860 Cryostat in 10µm sections and stored at -80°C until they were ready to be stained. COX-SDH staining protocol was adapted from Wanagat et al^34^. Briefly, heart sections placed on a slide were circled using a PAP pen, fully covered with COX-staining solution and incubated at 37°C for 12 minutes. Slides were then rinsed in 0.1M Tris-HCl and incubated at 37°C with SDH-staining solution for 10 minutes. Afterwards, slides were dehydrated in 70% ethanol, 95% ethanol, 100% ethanol (2x) and xylene before being coverslipped with Cytoseal 60. Slides were imaged on an Echo Revolve microscope at 4x, 10x, and 20x magnifications. Images were analyzed and scored for their % COX-negative cells using QuPath2 software.

### Statistics

Group differences means (± SEM) were analyzed by one-way ANOVA with Fisher’s LSD test, Kruskal-Wallis with Uncorrected Dunn’s test for nonparametric distributions, or Welch’s ANOVA when variances were unequal across groups. Significance was defined as p < 0.05. Analyses used GraphPad Prism version 10 (GraphPad Software, San Diego, CA).

## Acknowledgements

M.V. was supported by NIA awards R01GM124532, R01AG075130, and R01AG083065. S.R.K was supported by NIGMS grant R35GM133428. P.C. was supported by the Hevolution Foundation Award HF-AGE-23- 1273964-51. S.J.S. was supported by NIA fellowship F31AG084238.

## Author Contributions

MV designed the research. C.C. initially provided the animals necessary for the study. *In vivo* experiments were performed by SJS, EM, M.A, and IV. All other experiments were performed by S.J.S., E.M., L.C., S.L., H.A., C.S.C., G.D.L, B.M.V., J.W., and M.T. Bioinformatic analyses were performed by J.G.. M.V., S.J.S. and E.M. analyzed all data. M.V. and S.J.S. wrote the manuscript. M.S.C., P.C., S.R.K. edited and provided guidance for this manuscript.

---

**Figure S1.**
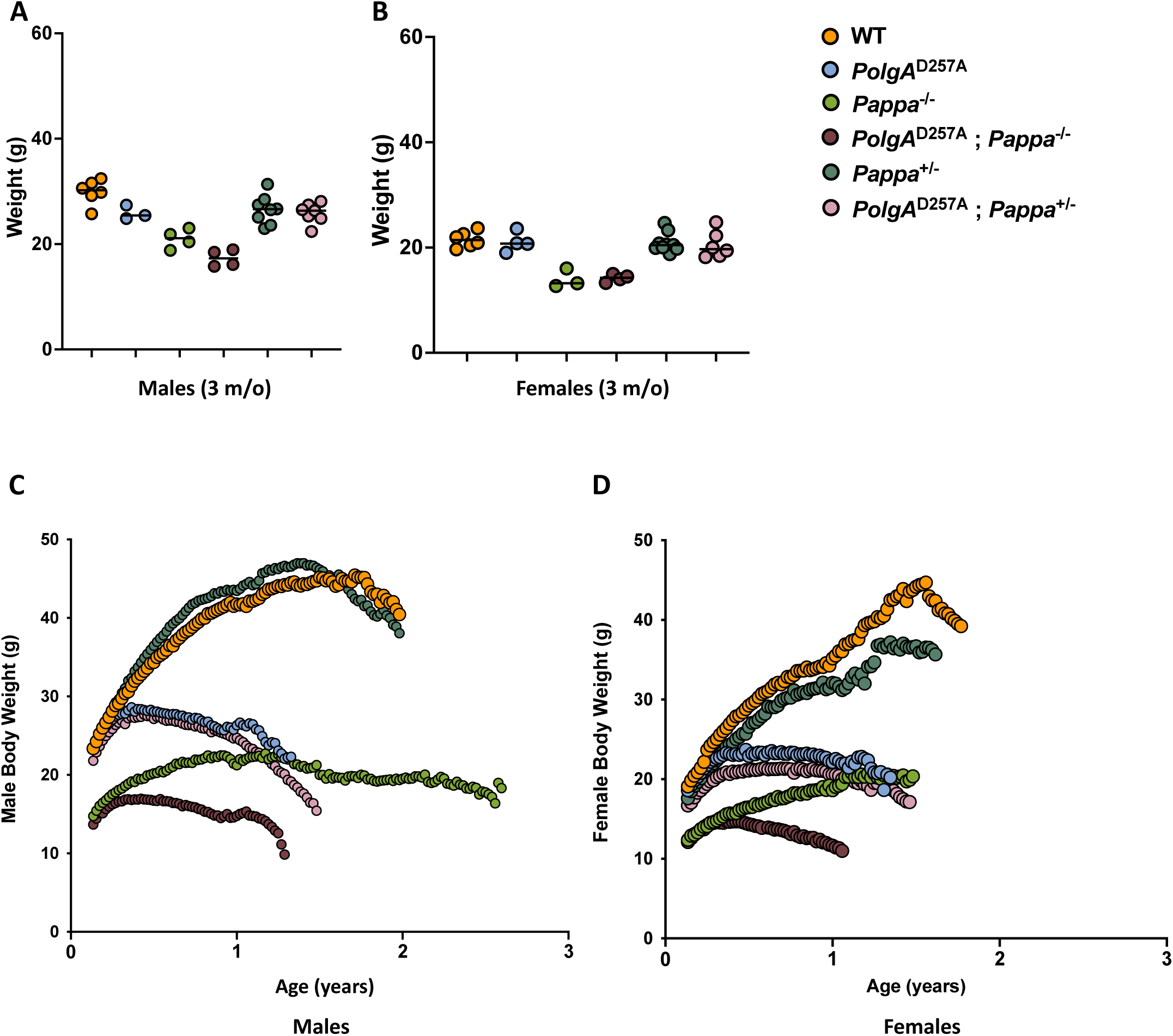
Weight distribution of young WT and mutant mice, and progression throughout their lifespan. **A.** Weight distribution of 3-month-old male mice (n=3-9/group). **B.** Weight distribution of 3-month-old female mice (n = 4-9/group). **C.** Weight distribution of male mice throughout their lifespan. **D.** Weight distribution of female mice throughout their lifespan.

**Figure S2.**
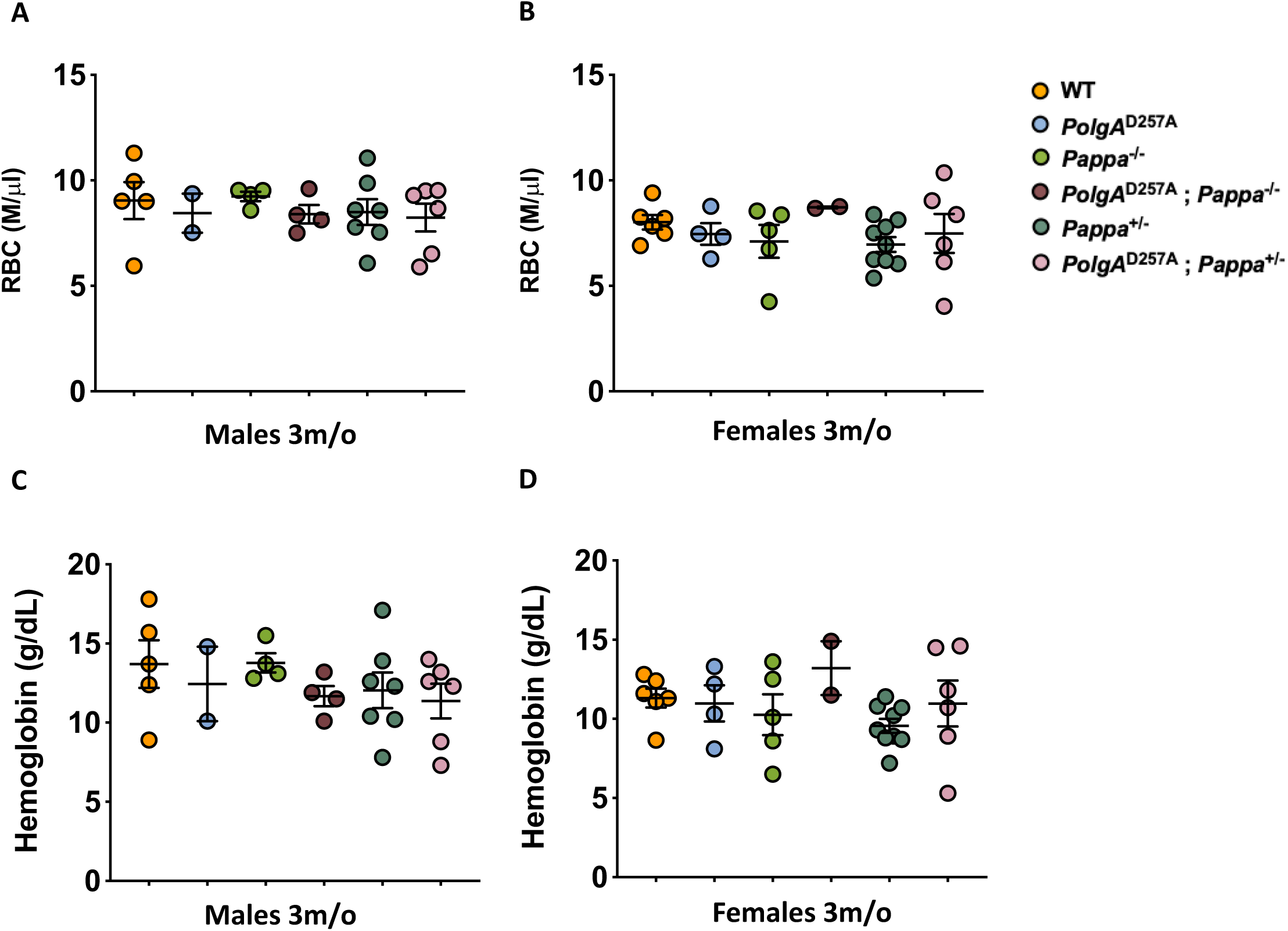
Red blood cell count and hemoglobin content in young WT and mutant mice. **A.** RBC count in 3-month-old male mice. **B.** RBC count in 3-month-old female mice. **C.** Hemoglobin content in 3-month-old male mice. **D.** Hemoglobin content in 3-month-old female mice. Male n = 2-7/group, female n = 2-9/group.

**Figure S3.**
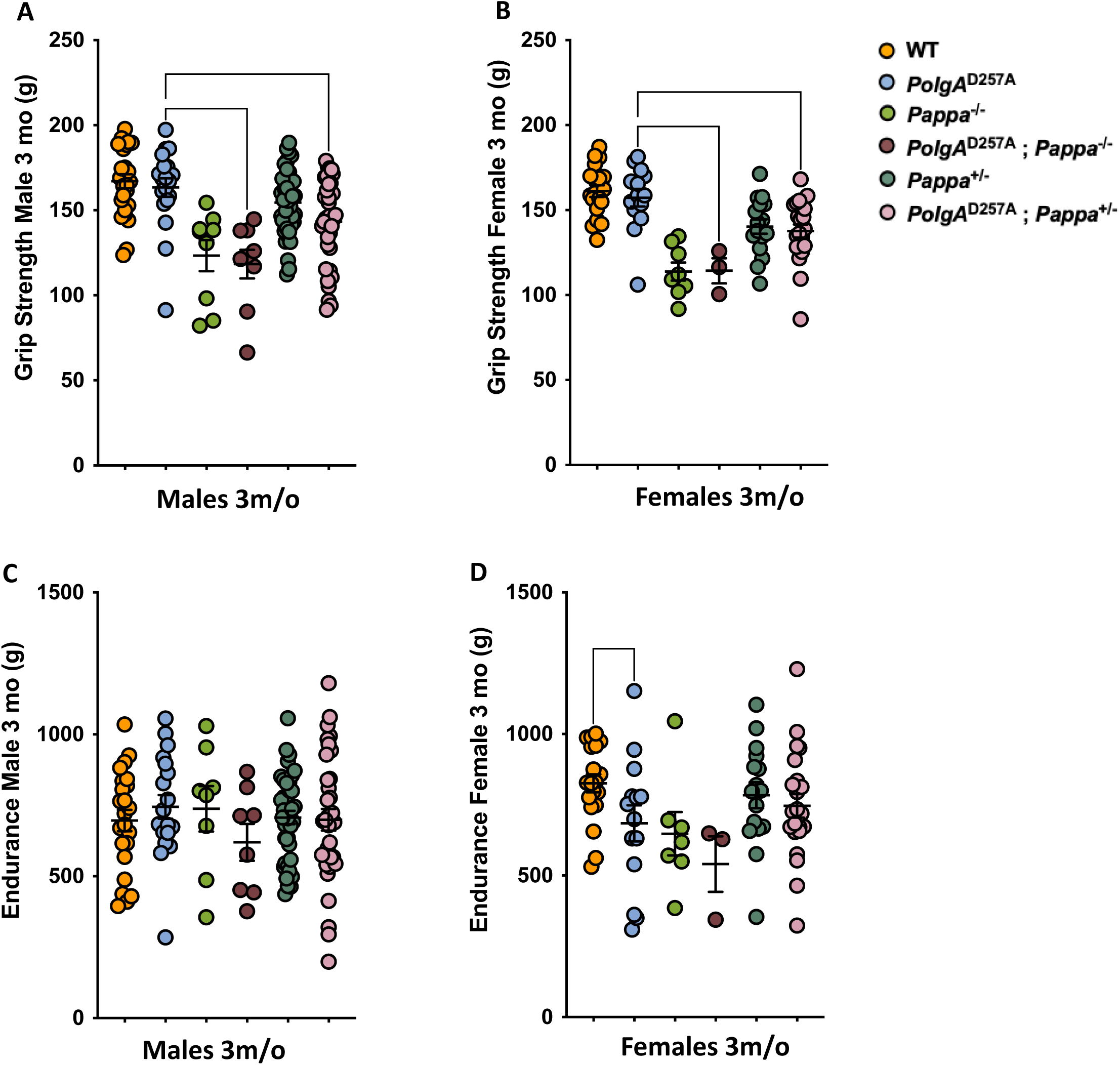
Grip strength and endurance in young WT and mutant mice. **A.** Grip strength in 3-month-old male mice. **B.** Grip strength in 3-month-old female mice. **C.** Endurance in 3-month-old male mice. **D.** Endurance in 3-month-old female mice. Male n = 9-40/group, female n = 3-22/group).

**Figure S4.**
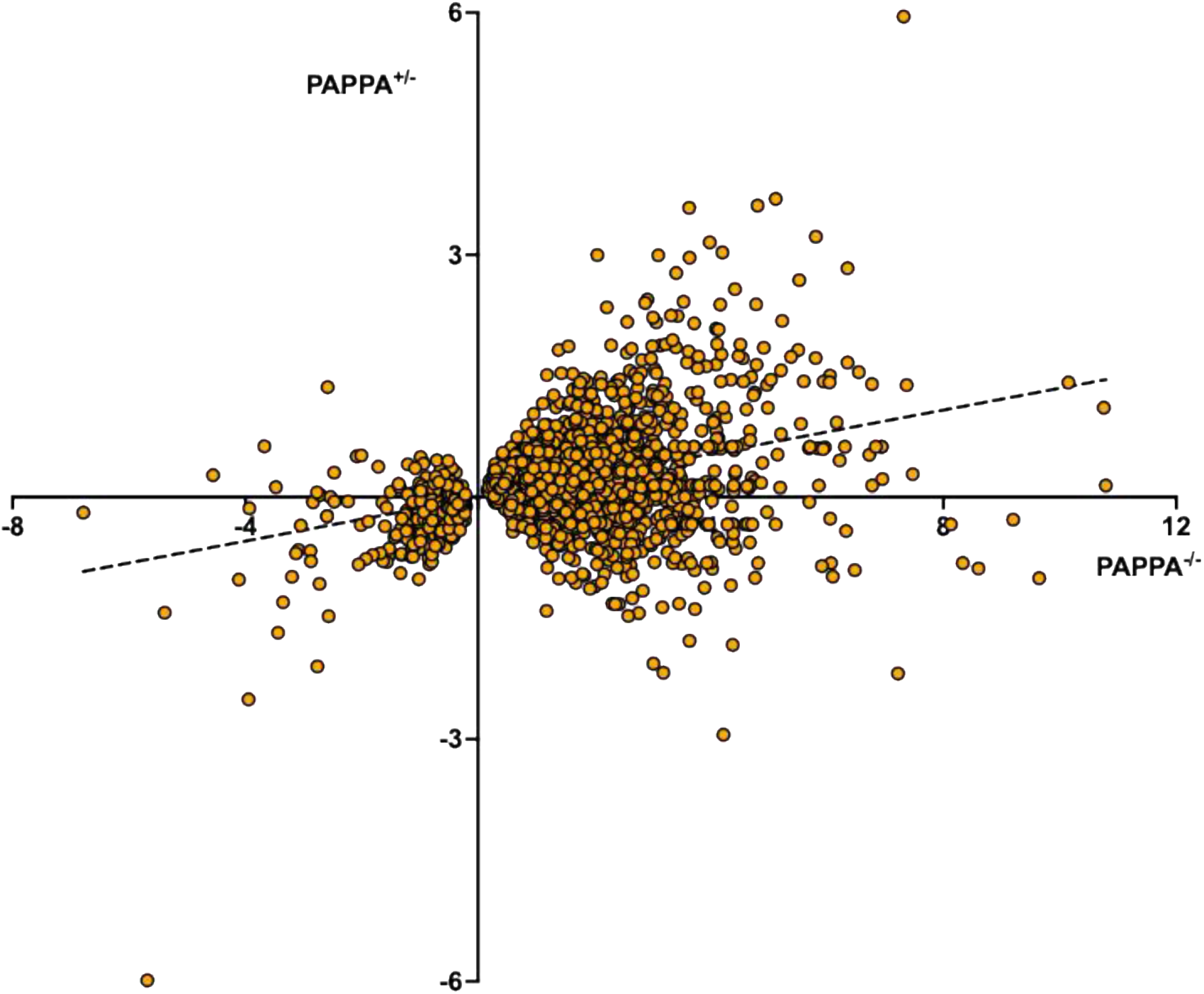
Gene expression in *Pappa*^+/-^ vs *Pappa*^-/-^ mice. Deletion of one copy of the *Pappa* gene shows a dose-dependent impact on gene expression compared to full deletion of *Pappa.* Genes that are upregulated in *Pappa*^-/-^ mice tend to be upregulated in *Pappa^+/-^* mice as well. Vice versa, genes that are downregulated in *Pappa*^- /-^ mice tend to be downregulated in *Pappa^+/-^* mice as well. However, the log2 fold change for each gene tends to be higher in *Pappa*^-/-^ mice vs *Pappa^+/-^* mice. Depicted are all genes that are significantly up or downregulated in *Pappa*^-/-^ mice. Both axes depict log2 fold changes, n = 6/group.

**Figure S5.**
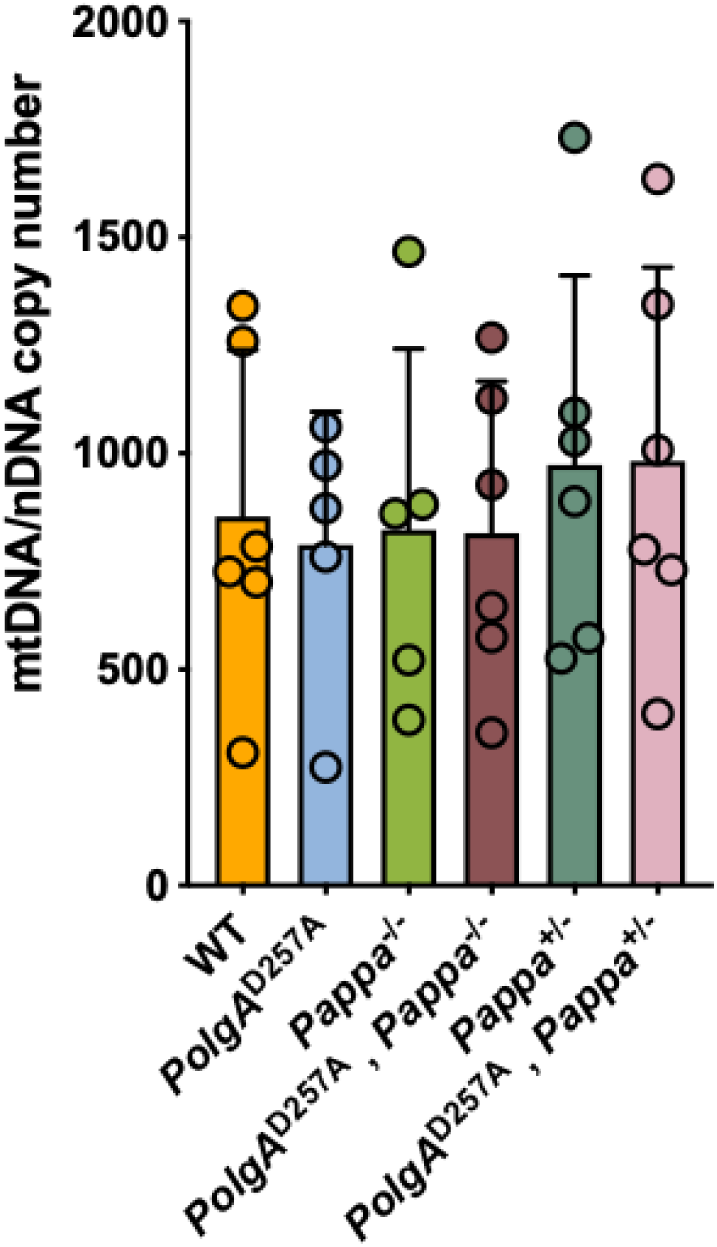
mtDNA copy number by qPCR in 12-month-old WT and mutant mice. qPCR detected no difference in mtDNA copy number between WT and mutant mice. N = 5-6/group.

**Figure S6.**
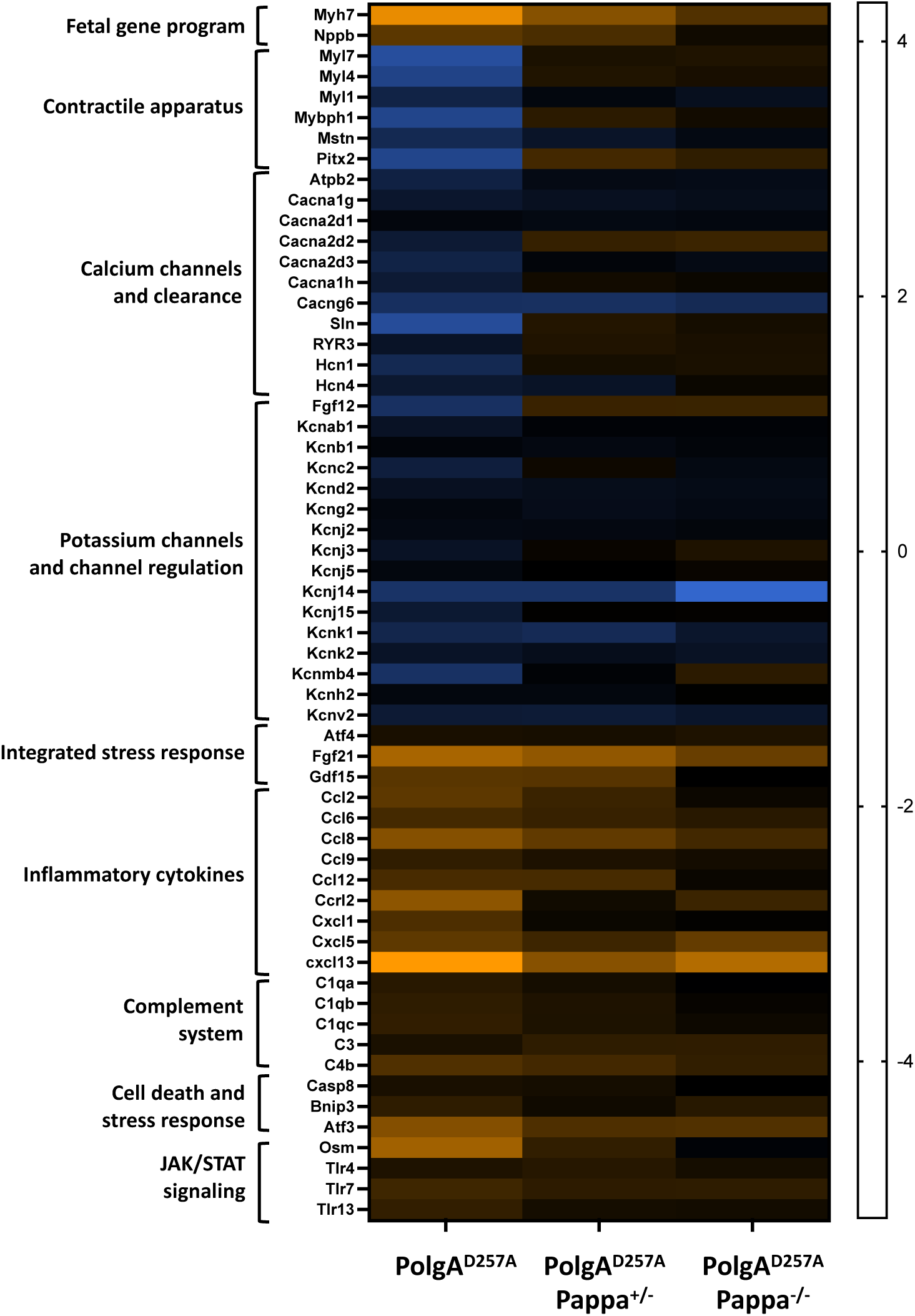
Expression of genes indicative of cardiomyopathy in male *PolgA*^D257A^ mice. Compared to WT mice, *PolgA*^D257A^ mice display multiple markers of cardiac stress, including increased expression of *Gdf15*, *Fgf21* and *Nppb*, as well as a reduction in the expression of genes key to contraction (*My1, Myl4 and Myl7*) and electrical signaling (calcium and potassium channels). Many of these genes move closer to WT levels upon deletion of one or two copies of the *Pappa* gene. *PolgA*^D257A^ mice also display increased levels of sterile inflammation, including inflammatory cytokines, components of the complement system, JAK/STAT signaling and markers for cell death. These inflammatory markers also seemingly subside upon deletion and depletion of the *Pappa* gene. N = 6/group.

**Figure S7.**
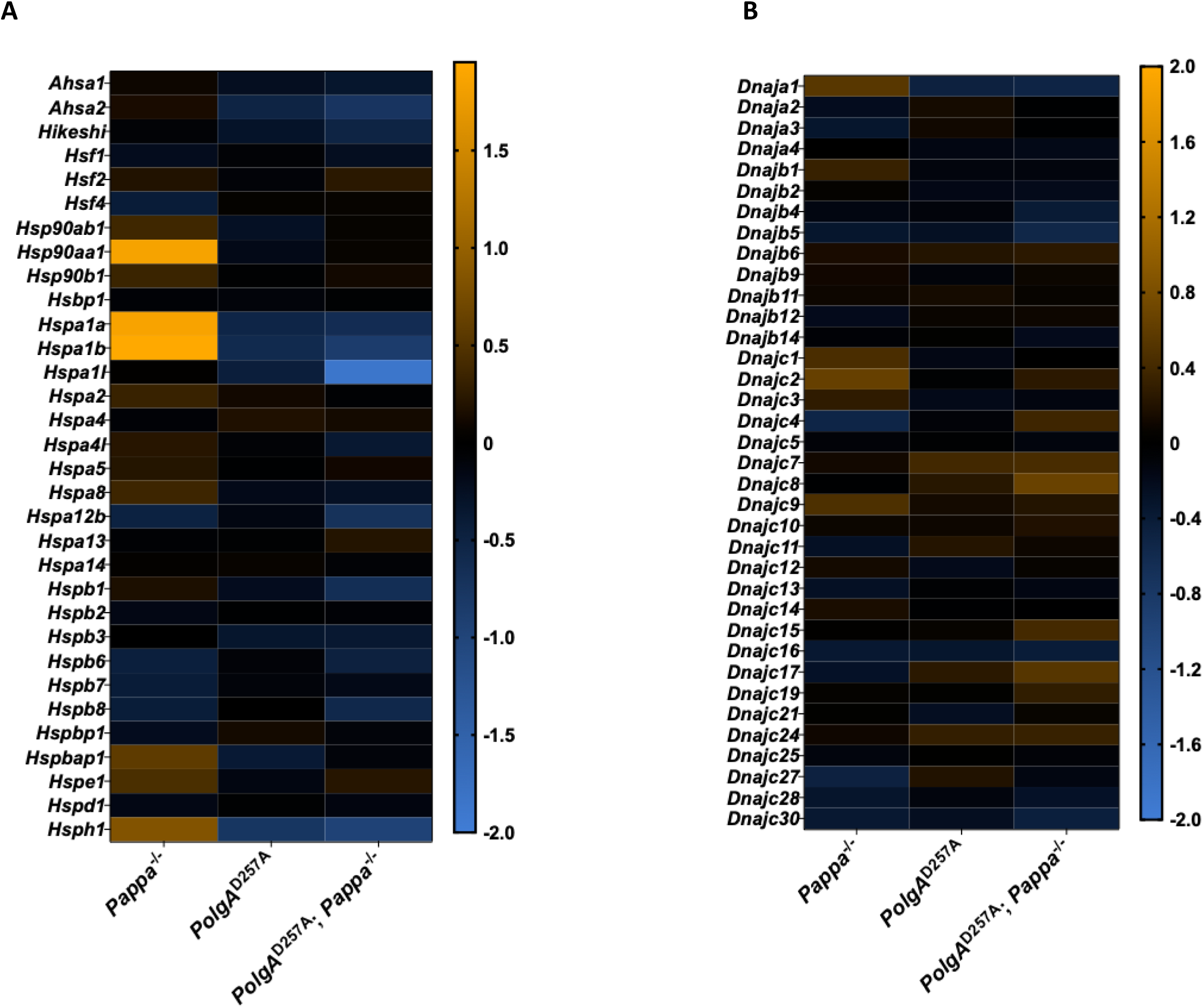
Expression of heat shock genes in *PolgA*^D257A^ mice with or without *Pappa* deletion. **A.** *PolgA*^D257A^ animals show downregulation of heat shock proteins, which is unable to be rescued by deletion of *Pappa*. These chaperones depend on ATP to function properly, stressing the importance of ATP production in their regulation. **B.** This is made clear by the loss of this trend observed in ATP-independent chaperone transcripts.

